# Cross-tissue patterns of DNA hypomethylation reveal genetically distinct histories of cell development

**DOI:** 10.1101/2022.12.15.520535

**Authors:** Timothy J. Scott, Tyler J. Hansen, Evonne McArthur, Emily Hodges

## Abstract

Establishment of DNA methylation (DNAme) patterns is essential for balanced multi-lineage cellular differentiation, but exactly how these patterns drive cellular phenotypes is unclear. While >80% of CpG sites are stably methylated, tens of thousands of discrete CpG loci form hypomethylated regions (HMRs). Because they lack DNAme, HMRs are considered transcriptionally permissive, but not all HMRs actively regulate genes. Unlike promoter HMRs, a subset of non-coding HMRs is cell-type specific and enriched for tissue specific gene regulatory functions. Our data further argues not only that HMR establishment is an important step in enforcing cell identity, but also that complex HMR patterns are functionally instructive to gene regulation. To understand the significance of non-coding HMRs, we systematically dissected HMR patterns across diverse human cell types and developmental timepoints, including embryonic, fetal, and adult tissues. Unsupervised clustering of 102,390 distinct HMRs revealed that levels of HMR specificity reflects a developmental hierarchy supported by enrichment of stage-specific transcription factors and gene ontologies. Using a pseudo-time course of development from embryonic stem cells to adult stem and mature hematopoietic cells, we find that most HMRs observed in differentiated cells (~70-75%) are established at early developmental stages and accumulate as development progresses. HMRs that arise during differentiation frequently (~35%) establish near existing HMRs (≤ 6kb away), leading to the formation of HMR clusters associated with stronger enhancer activity. Using SNP-based partitioned heritability from GWAS summary statistics across diverse traits and clinical lab values, we discovered that genetic contribution to trait heritability is enriched within HMRs. Moreover, the contribution of heritability to cell-relevant traits increases with both increasing developmental specificity and HMR clustering, supporting the role of distinct HMR subsets in regulating normal cell function. Altogether, our findings reveal that HMRs can predict cellular phenotypes by providing genetically distinct historical records of a cell’s journey through development.

**AUTHOR SUMMARY:** Studies aiming to understand the relationship between DNA methylation patterns and phenotypic outcomes have focused largely on individual differentially methylated regions without consideration of combinatorial changes that drive phenotypes. In non-disease contexts, most of the human genome is stably methylated, except for thousands of discrete DNA hypomethylated regions (HMRs) coinciding with gene regulatory elements. Here, we comprehensively characterize HMR relationships both within and between developmentally diverse cell types to understand the functional significance of complex HMR patterns. We show that levels of HMR specificity across cell-types captures time-point specific branchpoints of development. Our analysis further reveals that HMRs form clusters in proximity to cell identity genes and are associated with stronger gene enhancer activity. This is a wide-spread phenomenon and only a very small subset of HMR clusters is explained by overlapping super-enhancer annotations. Partitioned heritability revealed the functional significance of different HMR patterns linked to specific phenotypic outcomes and indicates a quantitative relationship between HMR patterns and complex trait heritability. Altogether, our findings reveal that HMRs can predict cellular phenotypes by providing genetically distinct historical records of a cell’s journey through development, ultimately providing novel insights into how DNA hypo-methylation mediates genome function.

## INTRODUCTION

Of twenty-eight million CpG sites in the human genome, the majority (80-85%) undergo constant DNA methylation (DNAme) (1–5). However, a subset of sites forms regions that are either broadly hypomethylated across cell-types or hypomethylated in a cell-type or lineage-dependent manner (hypomethylated regions, HMRs) (6–14). Contrary to long-held views of the role of DNAme in controlling promoter function, promoters are stably hypomethylated and largely invariant across cell types (15, 16). Thus, promoter HMRs poorly predict gene transcriptional status and ultimately cellular phenotypes. By contrast, the most dynamically hypomethylated HMRs are distal to promoters and often coincide with signature chromatin states typical of enhancers (5, 17–20). Yet, the functional significance of cell-type and lineage specific HMRs and their causal relationship with genes and cellular phenotypes is not well-defined.

Differential DNAme, like differential gene expression, is sufficient to distinguish diverse cell types and recapitulate their functional and developmental relationships (4, 9, 10, 21). Furthermore, differentially methylated regions are proximal to genes with tissue-specific functions and are enriched for GWAS (genome-wide association study) variants linked to tissue-relevant diseases (11, 22–25). While individual differentially methylated regions frequently overlap enhancers, few studies have considered the combinatorial significance of enhancer methylation patterns in a genome-wide manner across developmentally diverse datasets. For example, selective and continuous hypomethylation of individual enhancer units within super-enhancers (SEs) has been demonstrated in mouse embryonic stem cells (ESCs) during exit from naïve pluripotency, suggesting that coordinated hypomethylation of enhancers through cell fate transitions serves to uphold specific cellular states (26–28). Furthermore, our recent work shows that, while HMRs correlate with chromatin accessibility and other indicators of permissive chromatin, the temporal dynamics of HMR establishment are distinct from chromatin remodeling—HMRs persist long after chromatin remodeling changes (12). These studies indicate that HMRs uniquely capture both active and previously active gene regulatory elements; however, this hypothesis and its link to phenotypic outcomes remains to be tested across diverse tissues and developmental timepoints in a genome-wide manner.

Here, we performed a comparative analysis of whole genome methylation data from diverse tissues representing distinct organ systems and developmental timepoints. Unlike previous studies that emphasize pairwise differential methylation or locus-specific changes during limited differentiation time courses, we comprehensively characterize both HMR relationships within and between cell types to understand the functional significance of complex HMR patterns. We show that hierarchical conservation of HMRs across tissues can distinguish different types of enhancers and provides a map for broad developmental timelines underlying cellular states. We further demonstrate that this hierarchical relationship highlights meaningful HMR sequence contribution to complex trait heritability, ultimately providing novel insights into how DNAme mediates genome function.

## RESULTS

### Shared HMR patterns among diverse cell types reveal common functional and developmental histories

Non-coding HMRs can be functionally categorized by their cell-type specificity (9, 29). Previous work comparing distantly related cell types (embryonic stem cells versus terminally differentiated primary B cells) revealed two broad classes of HMRs: 1) highly shared, developmentally constitutive HMRs and 2) cell type specific HMRs that coincide with enhancer chromatin marks. Motivated by these studies, we hypothesized that differential comparisons of non-coding HMR profiles across multiple diverse cell types could provide insight into the presence and status of regulatory elements in individual cell types. To illustrate this idea, genome browser tracks of methylation data are displayed for datasets representing diverse lineages and developmental timepoints at a B cell enhancer cluster upstream of the CD27 gene (**Fig 1A**). A comparison of HMRs reveals different levels of cell-type and lineage specificity including HMRs conserved in all samples (developmentally constitutive), HMRs shared exclusively among lineage-related samples (e.g., hematopoietic cells), and HMRs present only in B cells.

**Figure 1:**
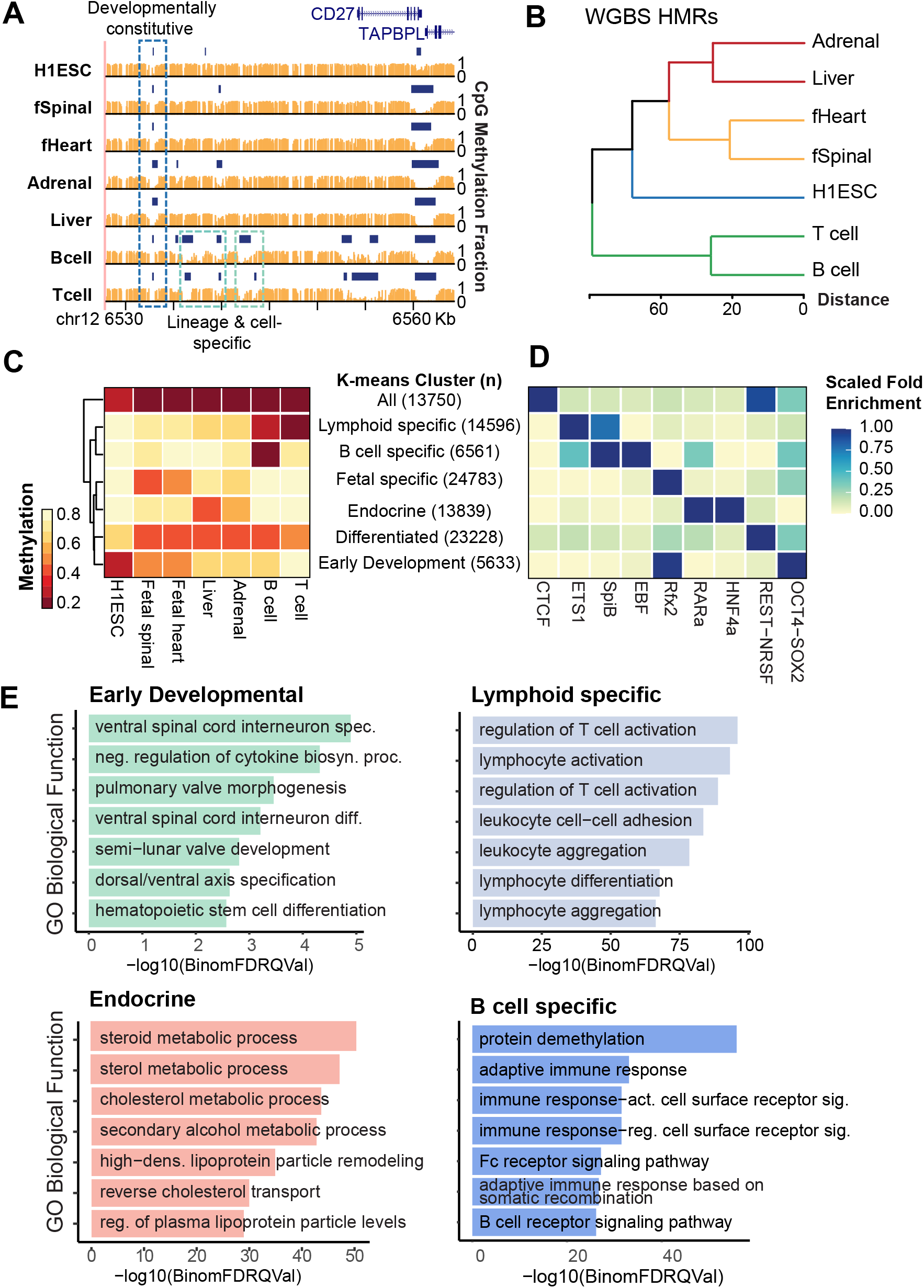
Levels of HMR specificity recapitulate developmental relationships. (A) Multiple alignment around the CD27 locus of methylation and HMR tracks across 7 cell types: H1 ESC, fetal spinal cord, fetal heart, adrenal gland, liver, B cell, and T cell. Methylation tracks are represented by orange vertical bars showing methylation value per CpG site. HMRs are shown by dark blue horizontal bars. Developmentally constitutive, lineage-specific, and cell-specific HMRs are highlighted. Sushi R package was used to generate genome browser snapshot (89). (B) Dendrogram of HMRs by average methylation per cell type through hierarchical clustering. (C) Heatmap of average methylation per HMR across cell types. Non-coding HMRs were k-means clustered based on their average CpG methylation values across 7 cell types represented in (A). A k-means of 7, assessed by the elbow method, was used to cluster HMRs into groups that are consistent with the biological relationships of their cell types. Groups are manually labeled to reflect their biological relationships. (D) The transcription factor (TF) motif enrichment of each k-means group reflects biological relationships captured in (A). Representative TFs were selected from top significant hits filtered by Benjamini-Hochberg adjusted p-value. Fold enrichment values are normalized from 0 to 1 across TFs. The background comparison file comprises HMRs across all represented cell types. (E) GREAT gene ontology enrichments are shown for selected groups from the heatmap in (C). Top 7 results by hypergeometric q-value are displayed. Bar plots show ontologies and q-values.

To investigate the intersection of HMR patterns globally, we determined a set of high-confidence HMRs using whole genome bisulfite sequencing (WGBS) data from 7 different cell types and tissues, including embryonic stem cells (H1ESCs), fetal heart, fetal spinal cord, liver, adrenal gland, T cells, and B cells (see Methods). As the resolution of HMR specificity is contingent on the quantity and interrelatedness of cell types included in the analysis, we maximized comparative potential by including datasets representing a diversity of organ systems and developmental stages; additionally, we used high quality methylation data with a minimum base pair coverage of >10x sequence reads (30). This resulted in a total set of 103,017 unique HMRs with an average length of ~784bp, excluding TSS and exon spanning HMRs.

Hierarchical clustering applied to these datasets was sufficient to recapitulate both related and distant cell type relationships, demonstrating the quality and specificity of HMR calls (**Fig 1B**). Next, we utilized *k*-means clustering to group non-coding HMR methylation levels across the 7 different cell types and tissues in an unsupervised manner. A gap statistic determined an optimal number of *k*-means clusters (n=7) and the resultant heatmap revealed groups of HMRs highly stratified by both cell type and developmental stage (**Fig 1C, S1 Fig**). We manually classified each *k*-means group according to cell types displaying average HMR methylation ≤50% for each group. For example, in the “Differentiated” HMR group all cell types except H1ESC display low methylation levels, whereas the “Early Developmental” HMR group is dominated by H1ESCs. Likewise, a group of HMRs is specific to the “Fetal” developmental state compared to stem and adult cells. Using this analysis, we achieve remarkable resolution to distinguish highly related cell types such as B and T cells; further, we identify a more specific group of exclusive B cell HMRs.

The specificity of the unsupervised groupings points to gene regulatory elements with shared functional and developmental histories. Since transcription factors (TFs) govern the functional progression and specialization of cell types, we performed TF motif enrichment analysis to understand the gene regulatory significance of each *k*-means group. Top motifs stratify strongly by *k*-means cluster (**Fig 1D**). Furthermore, representative TFs from top results show *k*-means cluster-specific enrichment of canonical TFs indicative of their respective cell types. For example, the ubiquitous CTCF is enriched in the *All* cluster (31); pluripotency factors OCT4-Sox2-Nanog are primarily in the *Early Developmental k*-means cluster (32); Lymphoid factor ETS1 is in the *Lymphoid specific* group (33, 34); and early B cell factor EBF is in the highly specific *B cell* group (35). Similarly, hepatocyte nuclear factor-4 alpha (HNF4α) and retinoic acid receptor alpha (RARα) motifs are present exclusively in the liver/adrenal-specific *Endocrine axis* group.

Given the specificity of the TF enrichment analysis supporting cell- and lineage-specific functions, we considered whether associated genes displayed similar biological specificity. Using GREAT ontology enrichment analysis (36), we show enrichment of distinct biological processes representative of the cell type and developmental stage associated with HMR groups (**Fig 1E**). Interestingly, T cell-related ontologies are strongly and uniquely enriched in the *Lymphoid specific k*-means cluster, whereas B cell-specific ontologies are enriched in the *B cell* group; this highlights the ability of HMRs to distinguish not only disparate lineages and developmental stages, but also highly related cell types. Together these data show that HMRs alone can recapitulate functional relationships between cell types. Furthermore, by comparing HMRs within and across lineages, we discovered that levels of HMR specificity can reflect deep developmental roots of gene regulation, capturing time-point specific branchpoints of development.

### HMRs accumulate and persist through subsequent developmental transitions

Terminally differentiated cells exhibit four times the number of non-coding HMRs compared to embryonic stem cells (**Fig 1C**). A minor subset of H1ESC HMRs are cell-type specific, while most H1ESC HMRs are highly shared across the cell types analyzed (2,822 vs. 15,413 non-coding HMRs). Our comparative analysis further reveals specific HMR groups defined by developmental stage (fetal vs. adult, differentiated vs. undifferentiated), lineage, and cell type (**Fig 1C**). These data suggest a model whereby H1ESCs supply a base HMR set to which additional HMRs are added at distinct lineage commitments through cell development. This is important because it suggests that a developmental hierarchy exists among HMRs and that HMRs accumulate as cells differentiate.

To determine whether progressive HMR establishment can be traced in developmentally derived cell types, we used publicly available WGBS data to construct a pseudo-time course of pluripotent (H1ESCs), multipotent (hematopoietic stem and progenitor cells, HSPCs), and terminally differentiated myeloid (macrophages) and lymphoid (B cells) lineage cells (**Fig 2A**). In general, we observe that HMRs increase in number with increasing cell maturity. An increase of total HMRs could be explained by (1) a simple accumulation of additional HMRs, or (2) a net increase with high turnover of HMRs. To differentiate between these two modes of HMR expansion, we measured HMR overlap between either embryonic stem cells or hematopoietic stem cells and mature hematopoietic cell types (**Fig 2B**). Of 18,235 HMRs observed in H1ESCs, 11,451 (62.79%) were represented in the multipotent HSPC dataset. In the fully differentiated cell types, 11,036 (60.52%) and 10,393 (56.99%) were maintained in the macrophage and B cell datasets, respectively. Next, of 34,605 HMRs established in HSPCs but absent in H1ESCs, 27,307 (78.91%) and 15,305 (44.23%) were represented in the macrophage and B cell datasets, respectively.

**Figure 2:**
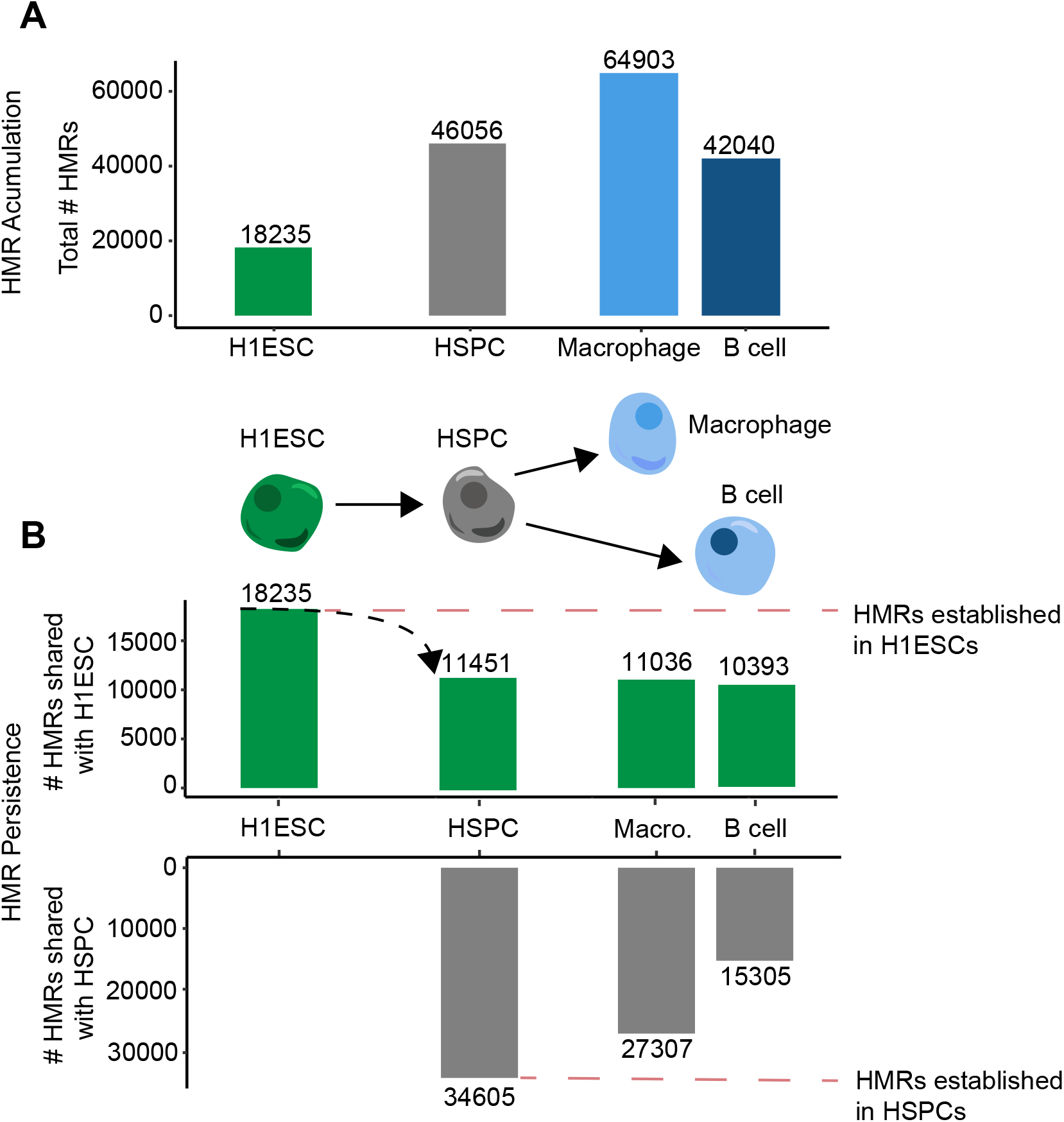
HMRs accumulate and persist through differentiation. (A) Bar graph of the total number HMRs for each cell type, arranged by developmental progression. (B) Bar graph measuring the presence of HMRs established in either H1 ESCs (*top; green*) or HSPCs (*bottom; grey*) in developmentally progressive cell types. The software Bedtools *intersect* was used to determine overlap between cell type HMR datasets using default settings. Overlap was defined as a 1bp minimum.

These data show that a majority of the HMRs observed in differentiated cells (~60%) are established at early developmental stages and suggest a pattern of HMR accumulation in relation to developmental progression. In addition to acquiring new HMRs, macrophages retain a majority of HMRs established in HSPCs, whereas B cells retain half as many HSPC-derived HMRs and fewer total HMRs compared to macrophages. This observation is consistent with previous studies demonstrating a lineage priming imbalance that favors myeloid differentiation in the absence of DNMT3A (37, 38); thus, lymphoid commitment and myeloid restriction requires re-methylation of specific early hematopoietic regulatory elements in parallel to demethylation of lymphoid-specific elements. Overall, our analysis supports a model where new HMRs are progressively established through successive developmental stages and persist through later stages of cell differentiation.

### HMRs are non-randomly established into spatially organized clusters

Locus-specific analysis of individual WGBS datasets indicates that multiple distinct HMRs are frequently located near one another, rather than being randomly distributed across linear genomic space. Moreover, these HMR groups appear to be spatially organized with HMRs displaying varying degrees of cell type specificity, from developmentally constitutive to B cell specific. An example locus shown in **Fig 1A** depicts a group of adjacent HMRs near the *CD27* gene, containing a cluster of genes which play a key role in B cell function (39–41). To quantify this genome-wide, we calculated observed and expected distributions of inter-HMR distances utilizing the broad set of diverse cell types represented in **Fig 1**. Expected distributions were simulated by random shuffling (n=10,000) HMRs across the genome for each dataset, excluding a blacklist of protein coding RefSeq promoters and exons. HMR distances are significantly closer to each other than expected by random chance (**Fig 3A,** Wilcoxon rank sum, p-value < 2.2e^-16^). Interestingly, differentiated cell types consistently show lower expected and observed distances (~40-50kb and ~12kb, respectively) compared to those of H1ESCs, which feature the largest inter-HMR distances; this is consistent with having fewer HMRs overall, supporting its role as a basal HMR set.

**Figure 3:**
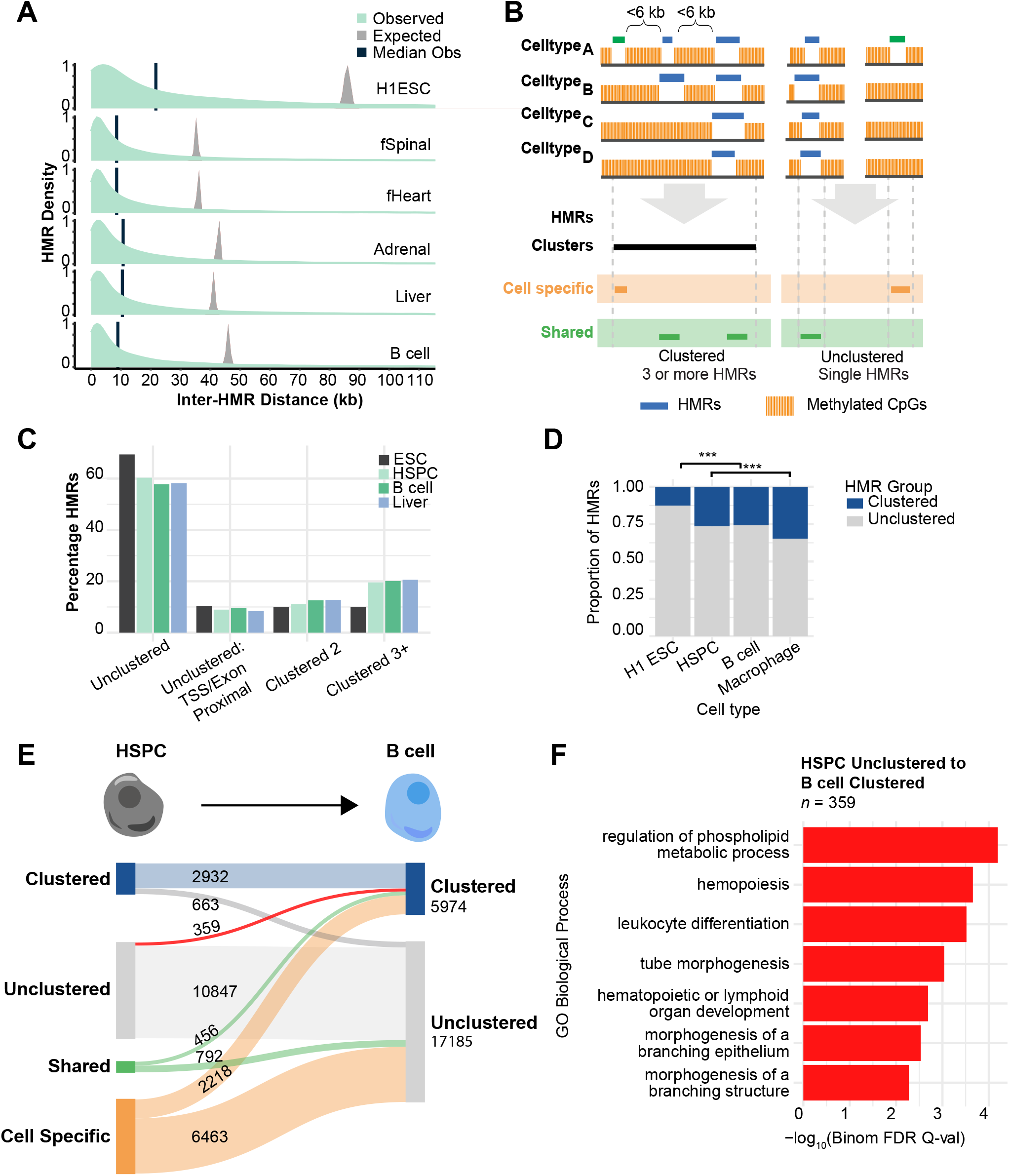
HMRs cluster more than expected. (A) Distribution plots of inter-HMR distances by cell type. The green distributions represent observed values from HMR datasets per cell type. Vertical black bars show median values. Gray distributions show expected values by random shuffling across the non-coding genome. (B) Diagram of HMR clustering and cell specificity workflow. HMR datasets are cleaned, removing regions that overlap RefSeq protein-coding promoters (TSS +2000/-1000) and exons. HMRs can be annotated for cell specificity and/or clustering behavior. Clustering refers to groups of HMRs in a cell type that are located a maximum of 6kb end-to-end from the next HMR, linking 3 or more HMRs. Cell specificity can also be annotated, with any base pair overlap between HMRs constituting overlap. (C) Bar graph of HMR clustering annotations discussed in (B) as percentages of total HMRs by cell type. Selected cell types represent members of the hematopoietic and hepatic lineages. Colors reflect cell types representing different developmental stages and lineages. (D) Bar graph of proportion of cell type HMRs that are clustered HMR. (E) Sankey diagram showing the flow of B cell HMRs. B cell HMRs are divided on the right of the panel into clustering groups. The left shows HSPC HMRs that overlap B cell HMRs, and are hierarchically categorized as *clustered HSPC* HMR, *unclustered HSPC HMR, shared*, or *cell specific*. To define cell specificity, B cell HMRs were compared to datasets from *adrenal gland, H1 ESC, HSPC, fetal spinal, fetal heart, liver, macrophage*, and *T cell*. (F) The bar graph shows the top biological process gene ontology results for the Sankey group of HMRs that progress from *HSPC unclustered* to *B cell clustered* (indicated in red). Results from GREAT Gene Ontology using default background and gene assignment settings are represented by bars showing binomial q-value.

These data suggest that clustered HMRs play a distinct regulatory role compared to their unclustered counterparts. To characterize the features that distinguish “clustered” and “unclustered” HMRs we first determined a set of heuristic criteria to define clusters (**Fig 3B**). We plotted per-cell type distributions for non-coding inter-HMR distances and measured distance quantiles. From this, we analyzed “end-to-end” cluster lengths based on three maximum linking values: the ≤12.5kb stitching distance commonly used in ChIP-seq-based super-enhancer studies (42–46), the approximate median HMR distance of ≤10.5kb, and ≤6kb which represents the median after filtering for end-to-end cluster lengths of 50kb (**S1 Table**). Previous studies that have characterized clustered super enhancers have used a common linking distance threshold of 12.5 kb. However, this distance was empirically identified in ChIP-seq datasets, and such a distance applied to methylation data results in extraneously long stitched regions, the biological function of which is difficult to assign; some exceed 1Mb, which can result from HMRs spread across gene deserts, or large topological domains with low methylation levels or CpG frequency. By comparison, a linking distance of 6kb results in stitched regions with an overall mean value of ~10.5 kb, which is consistent with other clustered enhancer annotations such as stretch and super enhancers ((43, 47), **S2 Table, S2 Fig)**. Using a linking distance of 6kb, we determined the fraction of HMRs that are clustered or unclustered for a subset of cell types, including H1ESC, HSPC, B cell and Liver (**Fig 3C**). To avoid confounding contributions of promoter characteristics to our analysis, clusters were not allowed to cross promoters or exons. At a 6kb threshold, non-promoter HMR groupings that exist as pairs or as clusters of 3 or more constitute ~35% of all HMRs.

As demonstrated in **Fig 1A**, a typical cluster consists of multiple HMRs with different levels of cell type specificity between them—broadly shared (developmentally constitutive), lineage-shared or cell-specific. This means that a cluster identified in one cell type, may not exist across all cell types, particularly in the case of H1ESCs. Moreover, many HMR clusters (~35-40%) contain at least one lineage- and/or cell type specific HMR. Given that HMRs accumulate over developmental timelines, this observation raises the possibility that HMRs are preferentially added to clusters in a lineage specific manner as cells differentiate. Indeed, we observe a positive correlation between clustering and developmental state. Clustering percentage increases as development progresses (**Fig 3D**, *H1ESC to HSPC* & *HSPC to Macrophage*: *p*<2.2E-16), and this is accompanied by a relative decrease in unclustered HMRs. Tracking these HMRs temporally for each pseudo-timepoint reveals that a substantial fraction of early-established HMRs is joined by additional HMRs in subsequent developmental states. The establishment of new HMRs near existing HMRs can lead to clustering, where an HMR may be classified as unclustered at an early timepoint but become clustered in a differentiated cell type. These growing clusters of HMRs are often in proximity to lineage-specific genes, as suggested by gene ontology analysis (**Fig 3E-F**). Altogether, these data show that HMRs can be broadly distinguished by 1) the number of cell types that share them – a corollary of temporal establishment or developmental time, and 2) their clustering behavior, which may reflect a collective and unique developmental function that distinguishes clustered HMRs from other types of genomic regions.

### Clustered HMRs are functionally distinct from unclustered HMRs

Sequencing approaches have enabled the discovery of many spatially clustered regulatory elements genome-wide using chromatin accessibility (47), histone modifications (48–53), and transcription factor binding (42, 43). More recently, clustering of enhancers has been commonly associated with concepts such as super enhancers (SEs) (43), stretch enhancers (47), shadow enhancers and locus control regions (LCR) (54, 55), which are thought to provide regulatory additivity, synergy, and redundancy to their target genes in a tissue specific manner.

Comparison of clustered B cell HMRs with histone H3K27ac-defined B cell super enhancers shows that, while HMRs capture a majority proportion of SEs, SEs explain only a minor proportion (~1.5%) of all clustered HMRs (**Fig 4A**). This discrepancy may be explained by the fact that SEs are often defined by strong H3K27 acetylation or strong TF binding, and only a fraction of SEs exist as clusters in linear genomic space. On the other hand, HMRs are more pervasive, and their clustering frequency may reflect a broader gene regulatory function or phenomenon. To address this question, we used ChromHMM annotations to functionally categorize HMRs based on clustering behavior in B cells (**Fig 4B**) (48, 49). Notably, clustered HMRs are enriched for “strong enhancers” (*X*^2^=46.358, *p*=9.848×10^-12^) while unclustered HMRs show higher enrichment of “heterochromatin” (*X*^2^=1434.3, *p*<2.2×10^-16^) and “insulators” (*X*^2^=731.47, *p*<2.2×10^-16^). This suggests clustered HMRs are enriched for active regulatory regions while unclustered HMRs tag elements involved in three-dimensional chromatin structure. This result is corroborated by the strong enrichment of the CTCF motif in unclustered HMRs, while both clustered and unclustered B cell HMRs show comparable enrichment of lymphoid-relevant transcription factors, including PU.1, SpiB, and ETS family members (**Fig 4C**).

**Figure 4:**
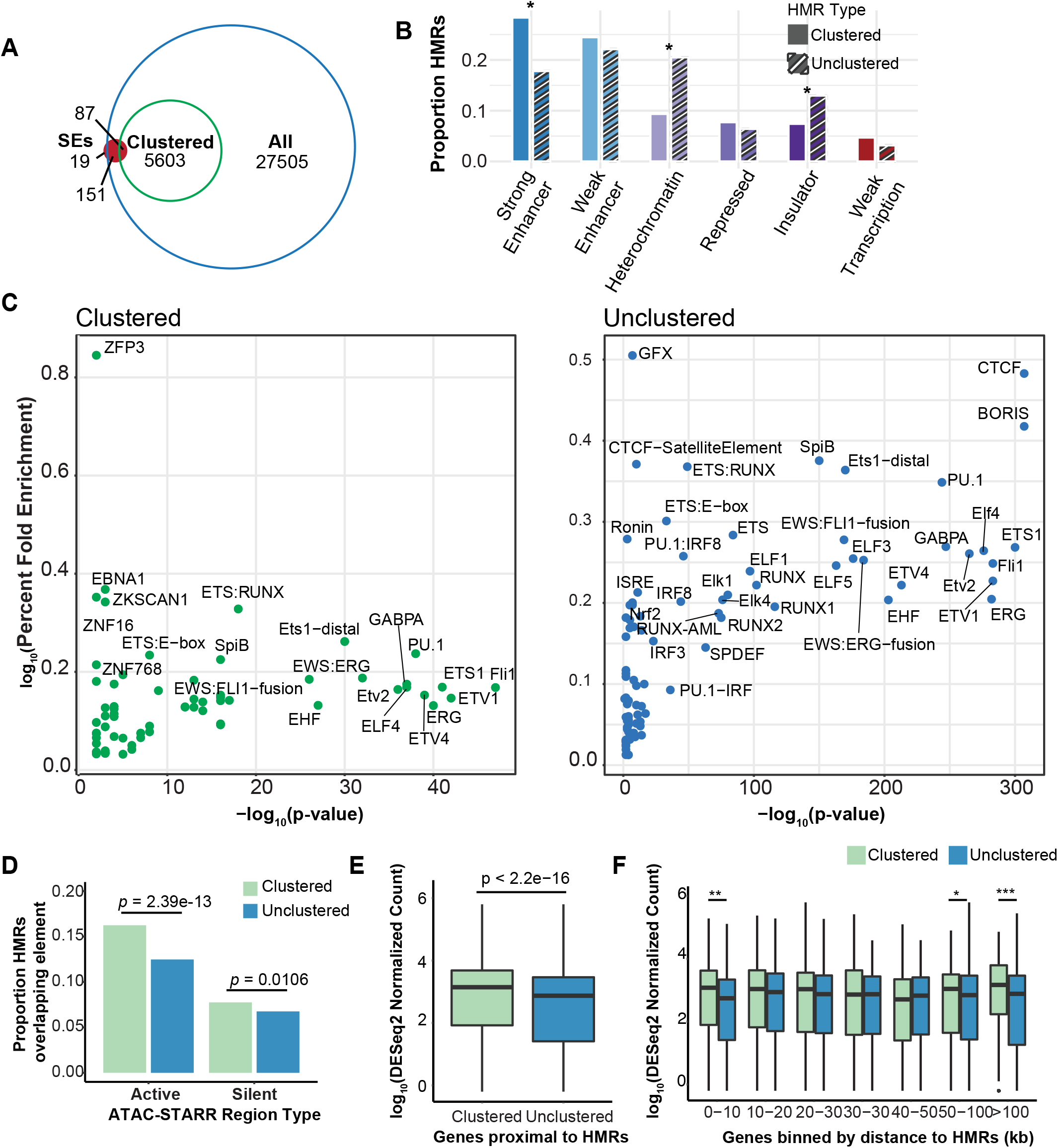
Clustered HMRs show distinct enhancer-associated characteristics compared to unclustered HMRs. (A) Venn diagram showing partially overlapping sets between three region datasets: All B cell HMRS (blue line); clustered B cell HMRs (green line; subset of All); and GM12878 super-enhancers (solid red circle) (42). (B) Bar graph of HMR overlap with selected ChromHMM annotations: *strong enhancer*, *weak enhancer*, *heterochromatin*, *repressed*, *insulator*, *weak transcription*. The height of the bars represents the fraction of clustered and unclustered HMRs that overlap each annotation. (C) TF motif enrichment in clustered (left; green) and unclustered (right; blue) HMRs. Results are plotted as −log_10_*p*-value by fold enrichment, measured as percentage of target regions containing motif divided by the percentage of background regions. Background represents all clustered and unclustered HMRs. (D) Boxplot of ATAC-STARR-seq regulatory element overlap by clustered and unclustered HMRs. Overlap is measured at the unit of HMRs, and values depict fraction of total HMRs that contain a regulatory element. A Wilcoxon rank sum test was used to determine statistical significance. (E) Boxplot of normalized read counts of nearest neighbor RefSeq protein-coding genes to clustered and unclustered HMRs. Nearest neighbor genes were filtered for TAD boundary crossing. Results are displayed for (E) all genes, or (F) genes binned by distance between the HMR and nearest gene. Statistical significance was measured by a Wilcoxon rank sum test (* = <0.05; ** = <5×10^-5^; *** = <5×10^-10^).

Given the enrichment of strong enhancer annotations in clustered HMRs, we investigated their transcriptional regulatory activity by comparing with our recently published ATAC-STARR-seq data for immortalized B cells (**Fig 4D**) (56). ATAC-STARR-seq is a massively parallel reporter assay that uses Tn5 transposase to selectively clone accessible DNA from native chromatin into a plasmid-based reporter to test accessible chromatin regions for active and silent regulatory activity (56, 57). Since a majority of B cell HMRs overlap accessible chromatin regions in lymphoblastoid cells (**S3A Fig**), we measured the proportion of HMRs that contain an activator or silencer (**Fig 4D**). Despite being fewer in number, clustered HMRs contain a significantly higher proportion of transcriptional regulators, including both activators and silencers (*p* = 2.39^-13^ and 0.0106, respectively), than unclustered HMRs.

Based on these results, we reasoned that target genes of clustered HMRs display increased transcriptional output compared to those of unclustered HMRs. To define putative HMR target genes, we used a gene assignment strategy that identifies the nearest expressed neighboring gene within a topologically associated domain (TAD) containing both the gene and the HMR(s). A recent study showed that a combination of nearest neighbor assignment in conjunction with a minimum expression threshold increased associated-gene prediction accuracy above several gene assignment methods, including the commonly utilized simple nearest neighbor (58). Using this assignment strategy and lymphoblastoid RNA-seq data from ENCODE, we observe that clustered B cell HMRs are associated with significantly increased gene expression (*p* < 2.2e-16; **Fig 4E**) (59). When these comparisons are binned by HMR-gene distance, significant differences in expression are observed at both proximal (<10kb) and distal (50-100kb) HMR-gene distances (**Fig 4F**). We performed the same analysis for liver clustered HMR target genes, finding the pattern is consistent across cell types (**S3B-C Fig**).

### Non-coding HMR patterns are highly enriched for genetic variants linked to specific clinical phenotypes

Genome-wide associations studies (GWAS) have demonstrated that a substantial portion of human phenotype-associated single nucleotide polymorphisms (SNPs) are located in functional regulatory elements (60–64). Integration of GWAS with functional genomic data reveals that disease risk variants also localize primarily within cell-type specific enhancers of disease-relevant tissues (65). Studies examining the relationships between disease loci and molecular phenotypes such as gene expression, chromatin accessibility or the DNAme status of *cis*-acting enhancers have identified a strong connection between non-coding genetic variants and epigenetic regulation (66–68). Based on these previous studies, we asked whether specific HMR patterns harbor genetic variants linked to cell-type-related clinical phenotypes, and, in turn, whether these relationships can inform the functional significance of different HMR patterns.

We reasoned that GWAS SNPS could be leveraged to reveal genetic variants in HMRs of critical importance to normal cell development and function. As in **Fig 2**, we defined B cell HMRs that are H1ESC-derived (developmentally constitutive), HSPC-derived (lineage-shared) or B cell-specific. We used stratified linkage disequilibrium score regression (S-LDSC) to perform partitioned heritability (69) analysis from GWAS summary statistics of 79 traits and clinical lab values representing a range of organ systems (**S3 Table**). We found significant enrichment of trait heritability within lineage- and cell-specific HMRs (**Fig 5A, S4 Fig**). Moreover, trait specificity enriches with increased HMR specificity. For example, H1ESC-derived B cell HMRs are nominally enriched for traits not immediately attributable to B cell function, such as *cardiomyopathy* and *morning person*. This is unsurprising due to the pleiotropy of gene regulation and the shared genetic architecture between many complex traits. However, HSPC-derived HMRs are enriched for genetic heritability of general hematopoietic traits including *white blood cell, platelet*, and *neutrophil counts*. In highly B cell specific HMRs, we identify a notable enrichment of specific immune-related clinical traits and lab values, several of which achieve significance after multiple testing correction (p<Bonferroni, n=79). These observations hold true for S-LDSC analysis in H1ESC-derived, and cell-specific liver HMRs (**S5 Fig**).

**Figure 5:**
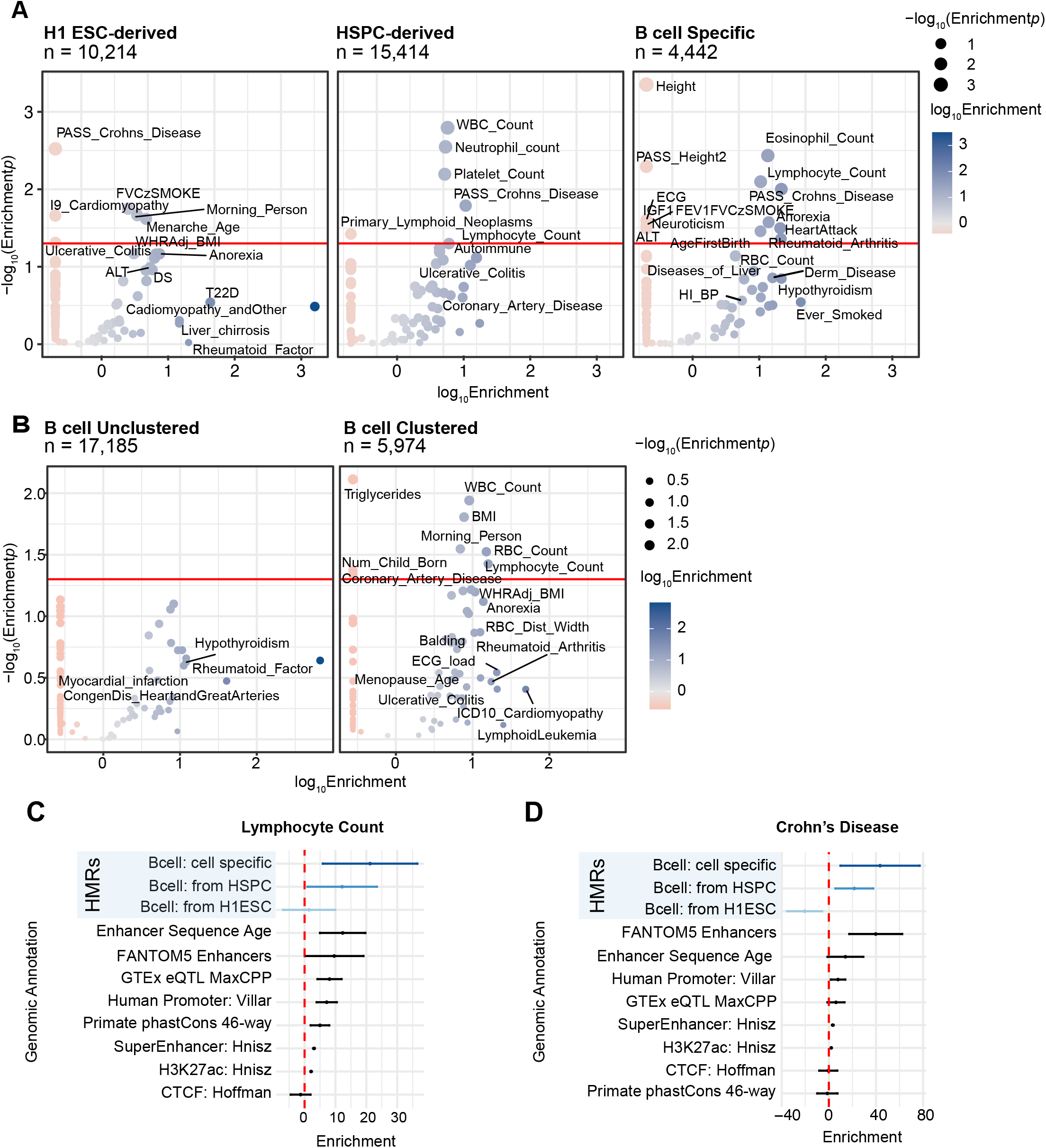
S-LDSC identifies HMR annotation-specific trait enrichments. (A) Volcano-style plots of S-LDSC partitioned heritability results across 79 traits are shown for three *B cell* HMR groups*: H1 ESC-derived, HSPC-derived, and cell-specific*. HMRs are ordered by the developmentally distinct cell type in which they were established. Each HMR group was tested for enrichment of genetic heritability with a standard set of 98 base annotations against traits that include both clinical diseases as well as clinical lab values. Negative enrichment values were clipped to the lowest positive enrichment value for each row of plots (A: 0.1174537; B: 0.25754925). The size of each point represents the −log_10_*p*-value of the enrichment, and the color shows the log_10_enrichment value. Points with a *p*-value <= 0.05 or an enrichment > 10 are labeled by their trait name where available. (B) Further partitioned heritability analysis applied to B cell HMRs grouped only by clustering behavior is also represented. (C) Point and line plots of S-LDSC enrichment values by annotation group per trait. These graphs include data from developmentally-derived B cell HMRs compared against other enhancer-associated groups, including ancient human enhancer sequence age, FANTOM 5 enhancers, eQTLs, super enhancers, and the H3K27ac histone mark. Genomic controls were also included, such as phastCons 46-way annotations as well as promoters and CTCF. The x-axis represents enrichment values, and the y-axis displays genomic annotations. Points show enrichment point estimates and lines display 95% confidence intervals. The red dotted line marks an enrichment score of 0.

HMRs stratified solely by clustering behavior also demonstrate heritability enrichment patterns associated with specific lymphoid traits (**Fig 5B**). In fact, compared to clustered HMRs, unclustered HMRs show no statistically significant trait enrichment above significance thresholds, suggesting that results in **Fig 5A** are powered predominantly by clustered HMRs. Accordingly, this trend was observed in gene-based disease enrichment analyses (disease ontology) applied to the same HMR groups analyzed by S-LDSC (**Fig S5**) (70, 71). For example, the top disease ontology enrichments for H1 ESC-derived HMRs uniquely include morphogenic ontologies such as *craniofacial abnormalities*, and the top ontologies for B cell specific HMRs include multiple lymphoid- and leukemia-related ontologies, reflecting the biological state associated with each HMR group. Together with the partitioned heritability results, these data suggest clustered cell-specific HMRs are both near lineage-specific genes and enrich for SNP variation associated with cell-specific traits.

To better contextualize the partitioned heritability enrichment results from B cell data, we compared results across cell types and against other known functional genomic feature annotations. We compared S-LDSC enrichment levels on a per-trait basis for B cell and liver HMR annotations (those from **Fig 5A and S4 Fig)** and other functional genomic annotations (**Fig 5C-D, S6A-B Fig**). For both immune-related clinical lab values and disease traits, we observe increasingly stronger enrichment from H1 ESC-derived to HSPC-derived to B cell-specific HMRs. In contrast, both H1 ESC-derived and liver-specific HMRs show positive enrichment for ALT (alanine transaminase) compared to B cell HMRs, as expected. This shows that SNP-based trait enrichment is capable of distinguishing HMR patterns from both distant and highly related cell types. Across cell relevant traits, we observe SNP-based heritability enrichment values that surpass those of promoters, expression quantitative loci (eQTLs), and histone marks of open chromatin (H3K27ac) often used to approximate active regulatory regions. Enrichment values associated with cell specific HMRs are comparable to those of FANTOM5 enhancers, supporting the notion that developmentally specific HMRs mark enhancers important for cell identity. Altogether, this analysis highlights the functional significance of different HMR patterns and indicates a quantitative relationship with complex trait heritability.

## DISCUSSION

While previous studies have characterized enhancer-associated methylation in a locus- or tissue-specific manner (72–75), we use comparative hypomethylation profiling to assess global hypomethylation patterns across cell types. This broader analysis reveals complex patterns of HMR establishment across a developmentally diverse dataset. By examining HMRs in a hematopoietic developmental context, we show that HMRs accumulate at distinct developmental stages and commonly persist through sequential lineage commitments. These developmentally hypomethylated regions are associated with distinct, stage-appropriate transcription factors, gene pathways, and SNP-based genetic heritability. These data allude to a model where H1 ESCs, with the fewest HMRs, present a basal set of HMRs to which additional regions are hypomethylated through development. In fact, about two-thirds of HMRs established in H1 ESCs remain in HMR datasets of differentiated cell types, highlighting their early establishment and continuous hypomethylation across time. Consequently, most (~3/4) HMRs in B cells were traced back to either H1 ESCs or HSPCs, indicating that the majority of HMRs are established at early cellular states. This further implies that biological differences between these cell types are driven by the minority population of differentially methylated HMRs. DMRs have been commonly used as a unit for studying DNA methylation (76–80); however, this annotation can lead to the loss of information of HMRs that are shared among various degrees of developmentally related cell types. Our results demonstrate that the entire HMR repertoire within a cell-type, rather than just the cell-type specific HMRs, stores information that is key to understanding and potentially predicting a cells phenotypic trajectory. In addition, we uncover a quantitative relationship between complex trait heritability and levels of HMR specificity, a feature that relates to HMR history. In general, we observe a strong pattern of hierarchical HMR establishment associated with stage-specific regulatory needs.

We further show that HMRs are preferentially established near existing HMRs, leading to the progressive enrichment of HMR clusters in differentiated cell types. Notably, clustered HMRs compose about 1/3 of all HMRs in differentiated cells, compared to less than 1/6 in H1 ESCs, indicating clustering increases proportionally to developmental progression. These spatially correlated HMRs are enriched for unique stage-relevant gene ontologies, trait-associated genetic heritability, and ChromHMM annotations, implying distinct regulatory roles compared to their unclustered counterparts. Previous investigations into enhancers describe subsets of clustering enhancers, including super-enhancers and hub enhancers (43, 54). While showing modest correlation with these annotations, HMRs capture additional non-coding space, suggesting clustered HMRs represent a broader epigenetic phenomenon than previously described.

We note that HMR clusters show patterns of hierarchical establishment that logically follow developmental paths. However, it is unclear if HMRs that persist through cell states remain epigenetically active at later stages. Clusters may include a combination of active and decommissioned, inactive enhancers recorded in HMR patterns. Murine models of early development have highlighted combinatorial and contrasting dynamics between spatially and functionally related enhancers during exit from pluripotency, where some require remethylation while others retain hypomethylation (27, 81). These enhancers may serve to uphold cellular states during cell fate transitions. Our lab has also observed ‘vestigial’ enhancers on shorter timescales by applying ATAC-Me-seq to a differentiation time course, simultaneously measuring chromatin accessibility and DNA methylation (82); these analyses reveal a subset of regions that undergo chromatin closing while simultaneously maintaining hypomethylation levels.

Partitioned heritability analysis of B cell HMRs established at three distinct developmental stages revealed enrichment of traits that reflect stage-relevant biology; in general, broadly shared HMRs were enriched for heritability of broader phenotypes while B cell specific HMRs were enriched for lymphocyte-relevant traits. Each B cell HMR subset likely suffers from power limitations, representing between 3,187,775 and 9,228,469 bp, or as low as ~0.1% of the genome. Despite this limitation, we observe a remarkable correspondence between heritability enrichment and stage-specific HMRs.

Our findings highlight DNA methylation as a unique epigenetic mark compared to common enhancer-associated histone marks. We highlight the unique accumulation of HMRs through developmental progression into clusters, enriched for stage-relevant SNP-based heritability. Through this process, epigenetic information can be maintained state to state. Thus, our results support that the methylome presents a historical documentation of developmental choices which could assist in the prioritization and interpretation of SNP data associated with clinical traits and diseases. These conclusions may further assist in understanding the complex role of the methylome in development and epigenetic gene regulation.

## METHODS

### HMR selection/exclusion dataset

DNA HMRs were obtained through the *MethBase* DNA Methylation trackhub from the UCSC Genome Browser, which references data processed through the *MethPipe* software for processing bisulfite sequencing data (30). To achieve a high-confidence genome-wide methylation dataset, cell types were included based on a minimum coverage of 30x (83). This resulted in the selection of: *adrenal, fetal heart, fetal spinal cord, liver*, and *T cell* from the NIH Roadmap Epigenomics Consortium (18); *H1 ESC* from Lister, et al. (4); and *B cell* from Hodges, et al. (9). As a primary cleaning step, with interest in only non-coding HMRs, we removed promoter- and exon-overlapping HMRs. To do this, we combined RefSeq exon and protein-coding gene TSSs (−1000, +2000 bp) annotations to form an exclusion BED file. Next, we referenced this file to eliminate promoter- and exon-overlapping HMRs using the *intersect* function from the *Bedtools* package with option ‘-v’ (84). Exclusion was defined by any basepair overlap. To increase the quality of our non-coding HMR dataset, we excluded HMRs based on a size minimum of 50bp. A subset of HMR datasets feature regions <8bp, suggesting artifacts from algorithmic noise, most likely incapable of featuring a functional TF binding site.

### HMR dendrogram

This analysis was performed using the per-HMR average CpG numerical matrix as composed in the methylation heatmap analysis. We used the *hclust(*) function and *ggdendro* package to perform hierarchical clustering and dendrogram composition in R.

### Methylation heatmap

Heatmaps were generated in R with the package *pheatmap (85*). Numerical matrices representing per-HMR DNA methylation per cell type were used as input. These were generated in bash using methylation bigWig files from the *MethBase* DNA Methylation trackhub hosted on the UCSC Genome Browser. We used the *KentUtils* binary package to convert bigWig files to bedGraph files. CpGs that likely mark CTCF binding sites were excluded, as measured by genomic overlap with a CTCF binding site as measured by Hoffman, et al. bedGraph score columns were used to populate a numerical matrix representing the sample-population methylation proportion at individual HMRs in rows. Heatmaps were generated using R package: *pheatmap* using options: *kmeans_k* = 7, *cluster_cols* = FALSE, *cuttree_rows* = 7.

### Transcription factor motif enrichment analysis

The *HOMER* perl package was used to calculate transcription factor motif enrichment(86). Background regions were defined per specific cell type dataset as cell type HMR-subtracted whole-genome hypomethylated space, represented by the merged HMR datasets of other cell types. Raw hypergeometric values were used to rank motif enrichment output. *Scale fold enrichment* was calculated by the quotient of two *HOMER* output values: *[%target/ %background]*. Top representative TF from the top 15 results are displayed in **Fig 1**. All data is represented in **Fig 4** to visualize TF enrichment differences attributable to clustering. Data visualization is scaled by TF to show relative cell specific enrichment. Graphing was performed in *R* using the ggplot2 package (87).

### *k*-means clustering gene ontology

Gene ontology was conducted using the web-based tool: *GREAT(36)*. Specifically, GREAT assigns regulatory domains around gene TSSs, extending to the nearest gene’s central domain up to a maximum extension distance. Here, we used the default gene annotation protocol from GREAT with a maximum extension of 1Mb. For a background file, we used a cumulative BED file representing all HMRs filtered for size and RefSeq promoters/exons from all cell types. Standard settings for maximum region-gene distance and gene assignment were used. Top results were downloaded from the web app using the “Shown ontology data as .tsv” selection. Output files were filtered to exclude the top row before import to R. Top ranked results are displayed as bar plots using ggplot2 and *geom_bar()*, ranked by binomial q-value.

### Inter-HMR distances

To measure inter-HMR distances, we employed the *Bedtools closest* function with the ‘-io -d’ options to calculate the distance from each HMR to the nearest HMR per cell type. Next, we extracted the output distance column to represent our observed distribution for graphing in R. To compare this distribution to random expectation, we used a script by Benton ML., 2018 which uses *Bedtools closest* to calculate distances between shuffled non-coding HMRs per cell type.

The *Bedtools closest* function takes two input files. For this analysis, the input dataset is submitted twice, though one input is shuffled. A region blacklist was used to exclude placement of HMRs during the shuffle into coding space, defined by RefSeq promoters and exons; this file was obtained from the HMR annotation step. We iterated the random shuffle-closest procedure 100,000 times to create a null expectation of genomic positioning given random placement. Distances per shuffle-closest were summarized as means, yielding a distribution of average distances per shuffle. Distributions were plotted using the ggplot2 R package.

### Clustering percentage

To assess the prevalence of clustering per cell type, we utilized the same procedures for annotating HMR clustering as were used in the primary HMR annotation script. Here, we specifically defined clusters by their amount of constituent HMRs; again, clusters were permitted a maximum of 6kb between HMRs. We used *Bedtools merge* with the ‘-d’ distance and ‘-c’ count flags to generate a dataset of merge regions. Regions that composed merged regions were then identified using *Bedtools intersect* with the original file as the -a file and merged datasets as -b files. By isolating various merged regions by number of constitutive HMRs, we were able to identify the quantity of HMRs involved in varying degrees of clustering. Data was compiled and binned into 1, 2, or 3+ HMRs. Graphing was performed with the ggplot2 R package.

### Sankey Diagram

HMR counts for each Sankey node and flow were determined using bash and the *Bedtools* suite. Nodes represent the total quantity of clustered and unclustered HMRs per cell type. Plots were generated in R using the package *networkD3*. To accurately represent the total quantity of HMRs per cell type, additional nodes were input and later processed with *Adobe Illustrator*.

### Sankey gene ontology

Gene ontology was conducted using the web-based tool: *GREAT(36)*, as with the *k*-means clustering gene ontology analysis. Here, we again used the default gene annotation protocol from GREAT. For a background file, we used the default “Whole genome” option. Standard settings for maximum region-gene distance and gene assignment were used. Top results were downloaded from the web app using the “Shown ontology data as .tsv” selection. Output files were filtered to exclude the top row before import to R, and the preceding “#” is removed from the second row. Top results as ranked by binomial q-value are displayed as a bar plot using ggplot2 and *geom_bar*.

### Super enhancer annotation

A super-enhancer catalog was downloaded from Whyte, et al. GM12878 SEs were isolated as a text file before comparison with clustered and all B cell HMR datasets using *Bedtools intersect*. Plotting was performed using the *R* package, *eulerr*, to maintain proportionally sized ellipses.

### ChromHMM annotation

A ChromHMM 15-state annotation file was downloaded from the UCSC Genome Browser in hg19 as assayed in the GM12878 cell line. Intersections were assessed using *Bedtools intersect* and B cell HMRs. Graphing was performed in *R* with the package, ggplot2.

### ATAC-STARR-seq comparison

BED files for GM12878 ATAC-STARR-seq regulatory elements were obtained from Hansen & Hodges (56) (GSE181317). HMRs were converted to GRCh38 for comparison using *liftOver* (parameters: -bedPlus=3). We determined the number of overlaps between the datasets with Bedtools *intersect* (default parameters) piped to a line count command (wc -l). The proportion of overlapping HMRs was calculated as [#overlapping/#total] and then plotted with *ggplot* in R. We performed a two-tailed, two-sample Z test of proportions with the *prop.test* function in R to obtain p-values.

### Nearest-neighbor transcriptional analysis

To measure the transcriptional differences associated with clustered or unclustered HMRs, we first identified a gene list with transcriptional values. To do this, we downloaded two replicate RNA-seq datasets acquired from the ENCODEv3 release. Here, we elected to use data for GM12878s, a lymphoblastoid cell line, to match B cells as closely as possible. Using the *DESeqDataSetFromHTSeqData, DESeq, collapseReplicates, and counts* functions with default options, we normalized replicate counts. Next, we combined *DESeq2* normalized count data with genomic coordinates, for comparison with HMRs in BED format, using the gene set defined by RefSeq *NM* genes. This file was downloaded from UCSC Genome Browser in *hg19* genome version and referenced earlier in the HMR annotation script. Gene coordinates and normalized counts were matched in R using the *merge* function. We also took this opportunity to add gene symbol identifiers with the *bitr* function from the *ClusterProfiler* package (version 3.0.4) for easier interpretability in downstream analyses. Next, to annotate nearest neighbor, we used *Bedtools closest* with the ‘-d’ distance option to identify associated genes with clustered and unclustered HMR separately. Using the previously described RefSeq promoter/exon file, *awk*, and *Bedtools intersect* (‘-v’ option), we filtered HMR-TSS pairs to keep those that do not cross a promoter or exon boundary. These were then additionally filtered against a GM12878 TAD file, downloaded from the 3D genome browser provided by the Yue lab (http://3dgenome.fsm.northwestern.edu/publications.html), to eliminate pairs that cross a TAD boundary. An expression filter (normalized count >0) was applied in R to eliminate HMR-TSS pairs that are likely inactive and a false positive pair assignment. The *tidyverse* package was used to bin HMR-TSS distances. ggplot2 and ggpubr were used for visualization in R.

### S-LDSC

Stratified LD-score regression was performed with *LDSC* using the appropriate python scripts distributed by the Price lab (https://github.com/bulik/ldsc). Reference base annotation files were downloaded from the Price repository (Phase 3, version 2.2 annotations). We used the appropriate reference files coordinating with the 1000 Genomes baseline v2.2 scores and HapMap 3 SNPs (https://alkesgroup.broadinstitute.org/LDSCORE/). Summary statistics were collected from both the Price lab (https://alkesgroup.broadinstitute.org/LDSCORE/independent_sumstats/) as well as the Neale lab heritability repository (https://nealelab.github.io/UKBB_ldsc/index.html). Traits were obtained based on their determined relevance to either broad cell-agnostic etiology or to biology specifically relating to B cells or Liver. This provided us the ability to determine specificity of results associated with varying cell specificity of HMRs. In total, we assessed 79 traits as described in **S3 Table**. The primary *LDSC* program was run per HMR annotation per trait. Results for individual traits were tabularized per HMR annotation. Results were visualized using *ggplot* in R with the functions *geom_point* and *case_when* for conditional coloring. To determine B cell developmentally derived HMRs, we used *Bedtools intersect* to compare HMR files. H1 ESC-derived B cell HMRs were defined by B cell HMR coordinates and had to had overlap with HMRs from HSPC as well as H1 ESC, together. HSPC-derived B cell HMRs had to have overlap with HSPC HMRs but not H1 ESC HMRs. B cell specific HMRs had to have no overlap with any HMRs from the collection of adrenal gland, liver, fetal heart, fetal spinal cord, H1 ESC, HSPC, Macrophage, and T cell. In the clustering analysis, all clustered or all unclustered HMRs were used. Liver HMRs were defined as H1 ESC-derived based on any overlap with H1 ESC HMRs. Cell-specific Liver HMRs were also defined against the same comparative cell type collection used with B cell for this specific analysis. Annotations used to compare against HMR groups were selected from those included in the “baselineLF_v2.2.UKB.tar.gz” from the Price lab LD-score website. Annotations were selected for their relevance to enhancers; CTCF, a ubiquitous transcription factor, was included as a negative control for cell-specific disease enrichment.

### WebGestalt Gene Ontology Analysis

Developmentally grouped B cell HMR BED files, as used in the S-LDSC analysis, were used as input in addition to BED files for all B cell clustered or all B cell unclustered HMRs. GREAT Input was used to identify nearest neighbor genes in hg19 (36). We used the default gene assignment parameters under “Association rule settings” called “Basal plus extension,” which in most cases replicates a two-nearest neighbor gene association strategy. In the web app, we downloaded the “Gene -> genomic region association table” file from the “genomic region-gene associations” page. Gene symbols were extracted from the GREAT input downloaded files using the first column. These were input into the WebGestalt web app to perform an over-representation analysis on the disease functional database, GLAD4U (70, 71, 88). For a reference gene set, we selected *genome protein-coding*. Results were downloaded, and the enrichment values file was used to plot enrichment ratio values for top diseases in R using ggplot2.

### NHGRI trait enrichment analysis

For SNP-associated trait enrichment analysis, we first obtained SNP-trait associations with the NHGRI SNP-trait catalog in *hg38* for information completeness (https://www.ebi.ac.uk/gwas/docs/file-downloads). HMR datasets were converted from *hg19* to *hg38* genome build versions using the *liftOver* tool from UCSC. *Bedtools intersect* was used to identify HMR-overlapping SNPs and associated traits. Resulting SNP-trait pairs were tabulated per HMR annotation group. The expected SNP-trait distribution was informed by the entire NHGRI catalog. Fold change was calculated against this null distribution. P-values were calculated using a binomial test to account for replacement and multiple SNPs associating with a single trait. Top results were ranked by p-value as displayed in the manuscript. Code was written in *Python* and graphing was performed in *R* using *ggplot*.

## ACKNOWLEDGEMENTS

We thank Tony Capra, Lea Davis, Melinda Aldrich, Eric Gamazon, Mary Philip, and members of the Hodges Lab for helpful feedback and discussions. We are grateful for support of the project and the time invested in producing this manuscript by NIH awards [1R01GM147078-01 to E.H, T32-GM080178-14 to T.S.], Department of Defense Idea Award [W81XWH-20-1-0522 to E.H], and American Cancer Society (ACS) Institutional Research Grant (#IRG-15-169-56).

## SUPPLEMENTARY FIGURE LEGENDS

**S1 Fig.**
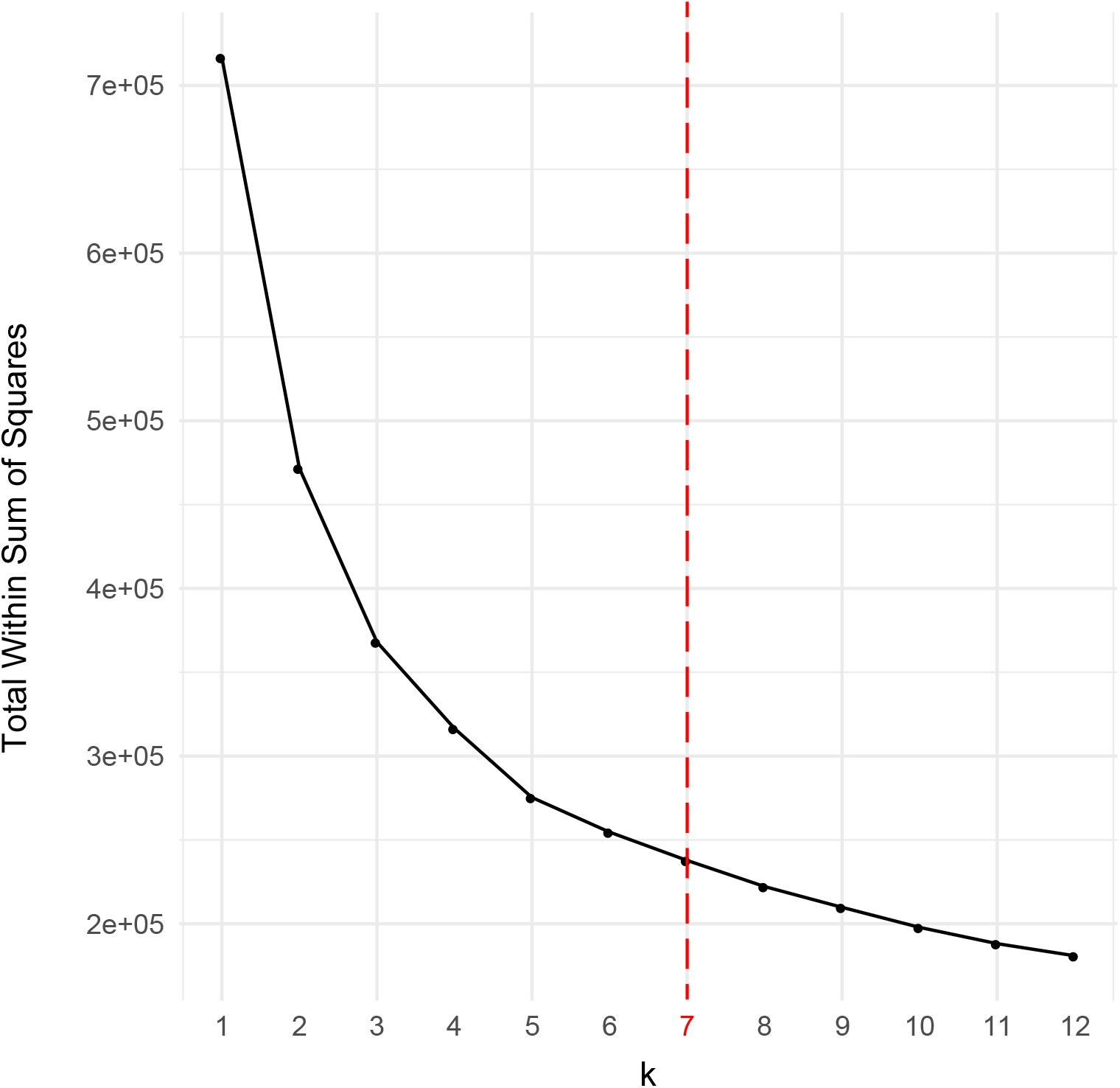
Dotplot of elbow method to determine appropriate number of k-means for methylation heatmap. Figure displays within sum of squares estimates for clusters at each value of k-means group amount from 1 to 12. Estimates are derived from the *kmeans(*) function in R.

**S2 Fig.**
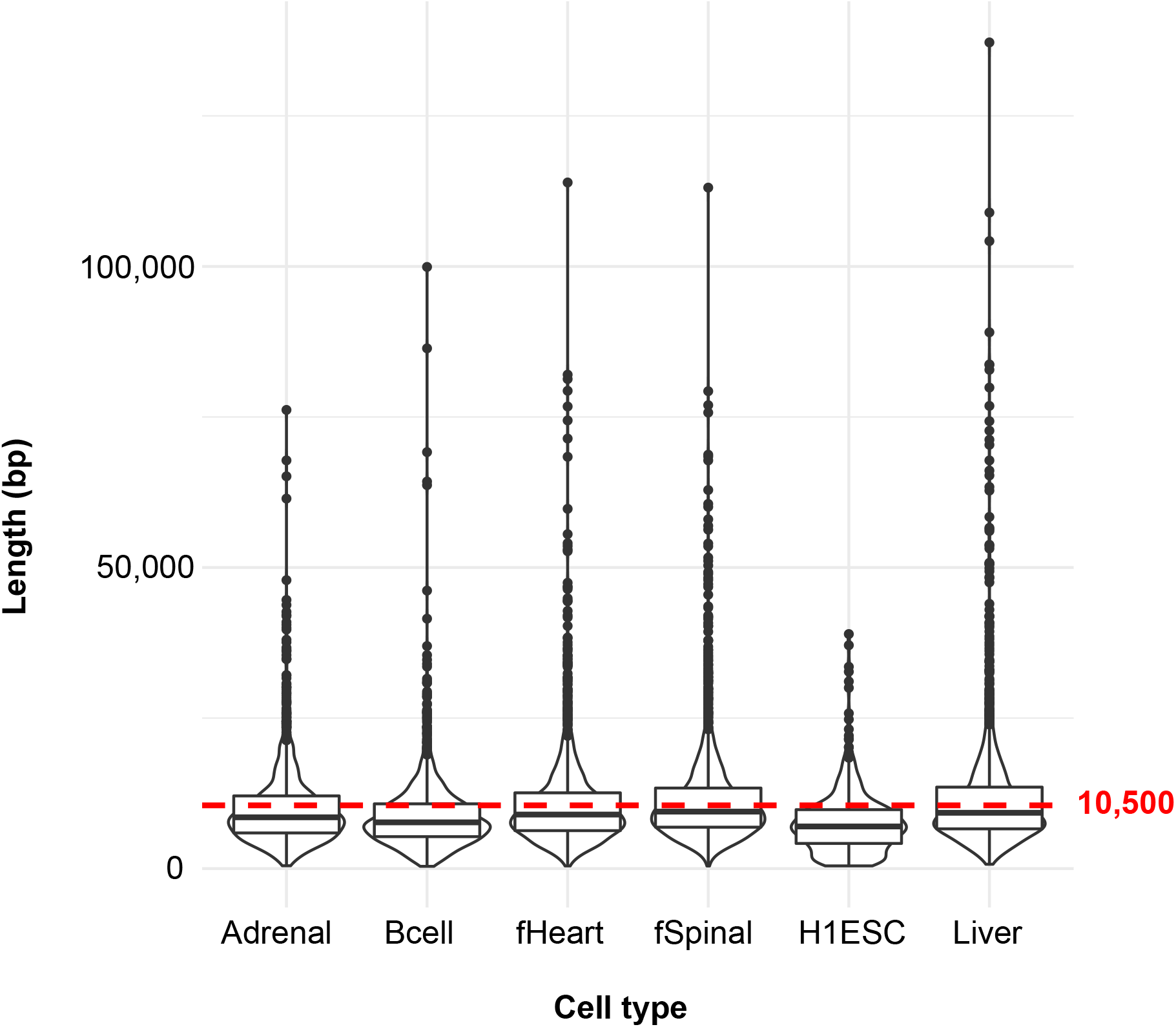
HMR cluster lengths are consistent across cell types. The graph shows the lengths of HMR clusters, end-to-end, per cell type. Data is represented by both a violin plot and boxplot. The boxplot shows the interquartile range, and the bold black line shows the median value per cell type. The red dotted line shows the value 10,500 bp, which approximates the mean cluster length of 10,551.2 bp, measured across the cell types: *adrenal gland, fetal heart, fetal spinal cord, H1 ESC, and Liver*.

**S3 Fig.**
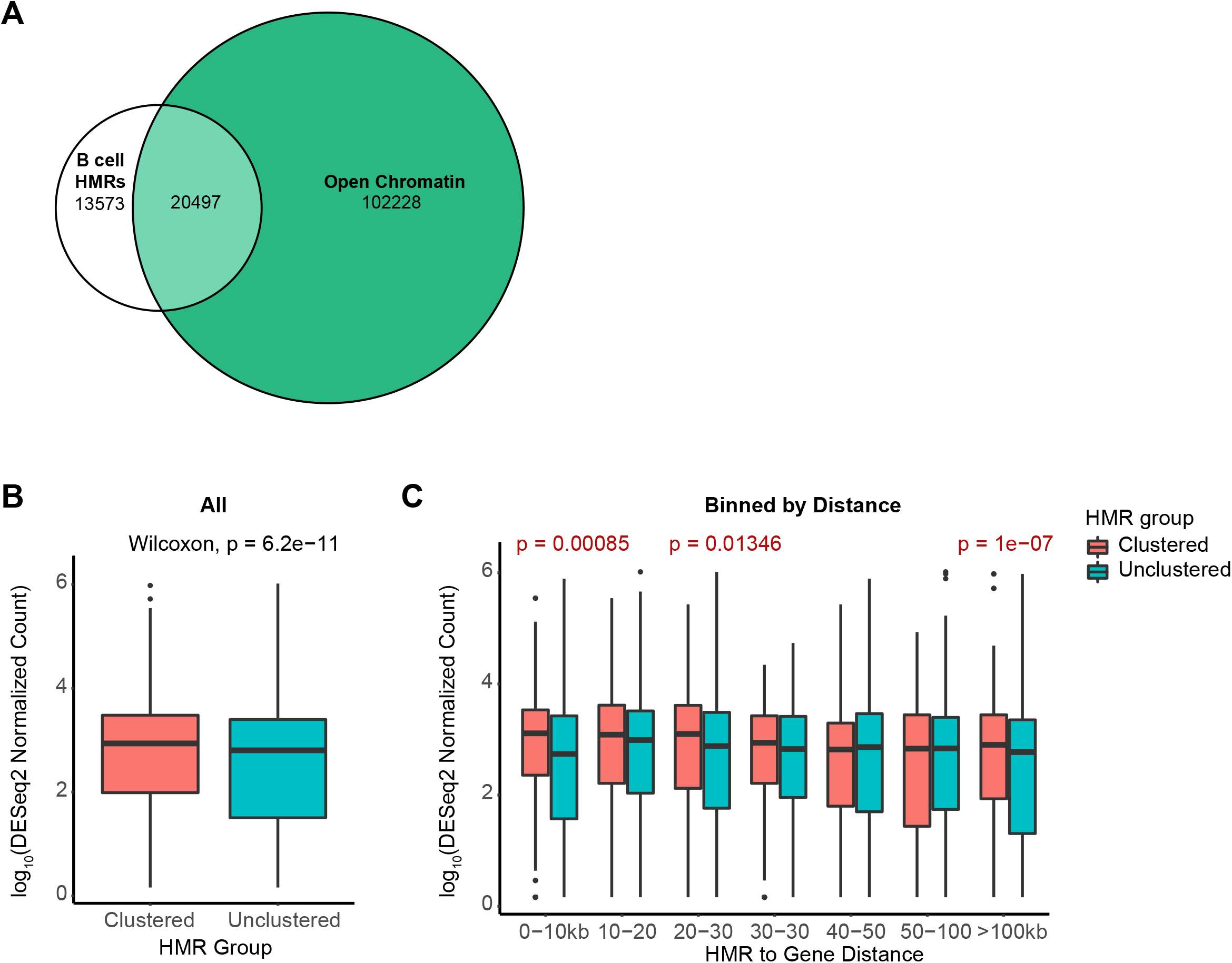
Euler plot comparing B cell HMRs and open chromatin. (A) Euler plot of all B cell HMRs and open chromatin defined by DNase I hypersensitivity sites in GM12878 cells. The DNase file was downloaded from the UCSC Genome Browser Table Browser using the following main settings: *clade*: “mammal”; *genome*: “human”; *assembly*: “Feb 2009 (GRCh37/hg19)”, *group*: “Regulation”, *track*: “Duke DNasel HS”, *table*: “GM12878 Pk (wgEncodeOpenChromDnaseGm12878Pk)” [ENCODE file ID: ENCFF001UVC]. (B-C) Boxplot of normalized read counts of nearest neighbor RefSeq protein-coding genes to clustered and unclustered Liver HMRs. Nearest neighbor genes were filtered for TAD boundary crossing. Results are displayed for (B) all genes, or (C) genes binned by distance between the HMR and nearest gene. Statistical significance was measured by a Wilcoxon rank sum test (* = <0.05; ** = <5×10-5; *** = <5×10^-10^).

**S4 Fig.**
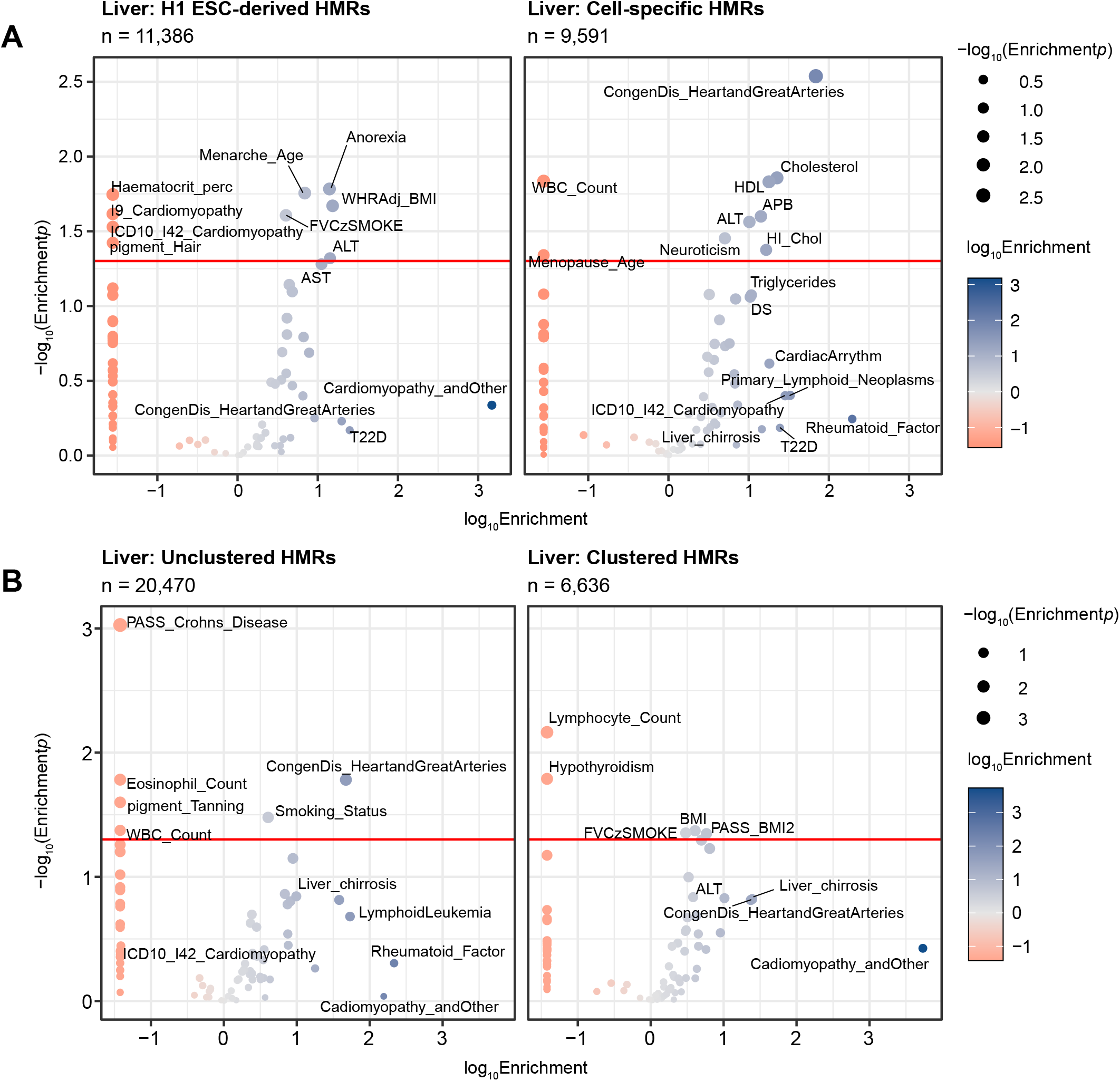
S-LDSC identifies HMR annotation-specific trait enrichments in Liver. (A) Volcano-style plots of S-LDSC partitioned heritability results across 79 traits are shown for two *Liver* HMR groups*: H1 ESC-derived and cell-specific*. HMRs are ordered by the developmentally distinct cell type in which they were established. Each HMR group was tested for enrichment of genetic heritability with a standard set of 98 base annotations against traits that include both clinical diseases as well as clinical lab values. Negative enrichment values were clipped to the lowest positive enrichment value for each row of plots (A: 0.02781896; B: 0.03787533). The size of each point represents the −log_10_*p*-value of the enrichment, and the color shows the log_10_enrichment value. Points with a *p*-value <= 0.05 or an enrichment > 10 are labeled by their trait name where available.

**S5 Fig.**
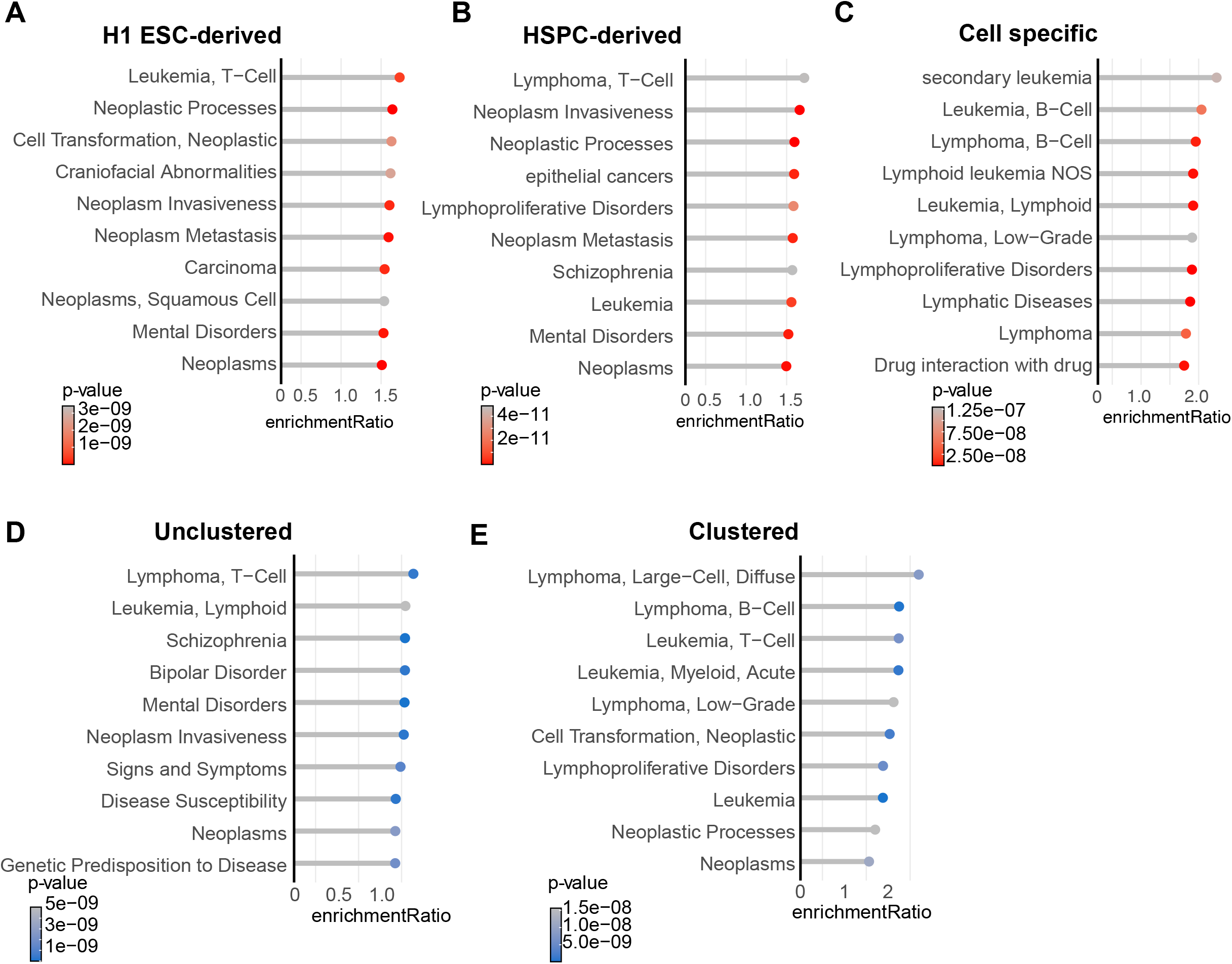
Disease ontology for developmentally specific and clustered B cell HMRs. Lollipop plots show top ten disease ontology enrichments as analyzed through WebGestalt with default parameters. The x-axis shows enrichment ratios, and the y-axis displays disease ontologies sourced from the GLAD4U disease database. The y-axis is sorted by enrichment value. The color for each bar represents the p-value for that trait. Individual graphs show results from B cell HMR developmental and clustering groups: (A) *H1 ESC*-derived, (B) *HSPC*-derived, (C) cell specific, (D) clustered and (E) unclustered.

**S6 Fig.**
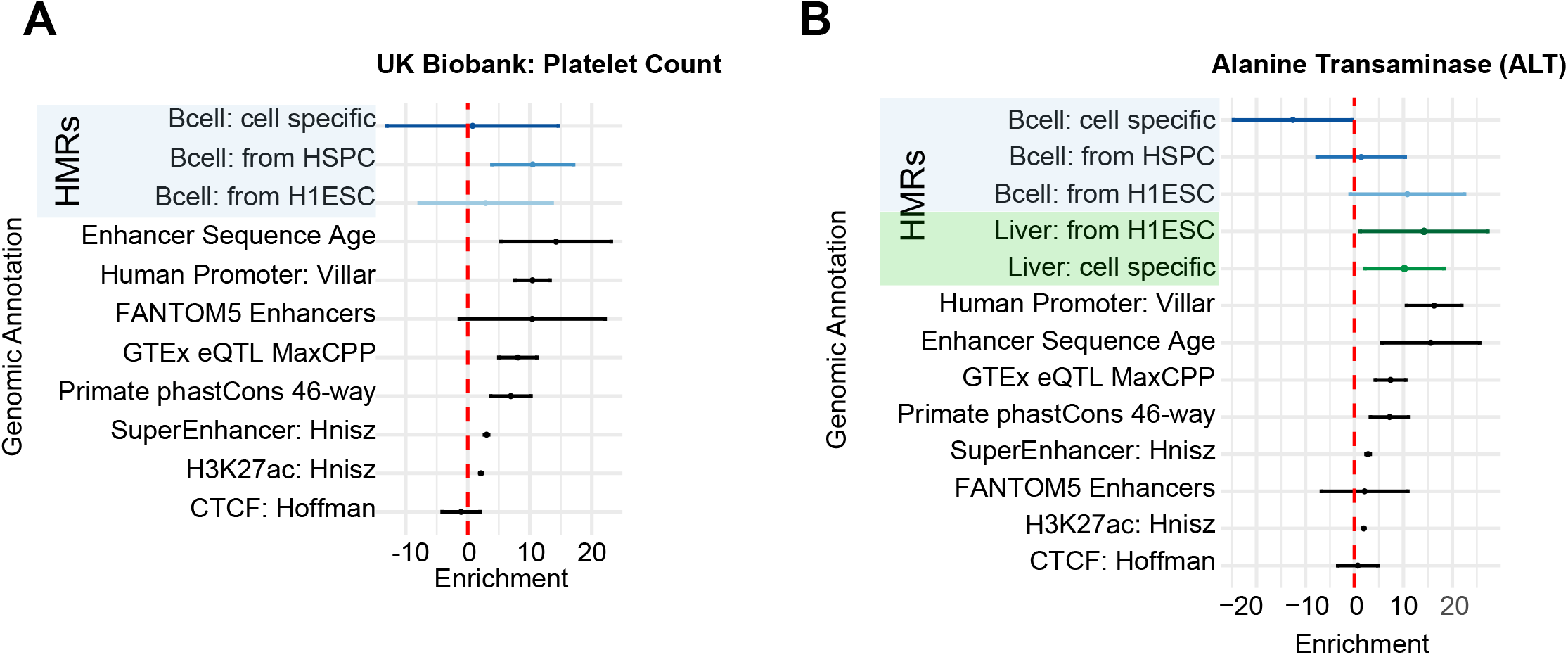
S-LDSC B cell by trait across genomic annotations. Point and line plots of S-LDSC enrichment values by annotation group per trait. The x-axis represents enrichment values, and the y-axis displays genomic annotations. Points show enrichment point estimates and lines display 95% confidence intervals. The red dotted line marks an enrichment score of 0. Annotation groups include popular enhancer-associated groups such as ancient human enhancer sequence age, FANTOM 5 enhancers, eQTLs, super enhancers, and the H3K27ac histone mark. Genomic controls were also included, such as phastCons 46-way annotations as well as promoters and CTCF. These graphs include data from (A) developmentally derived B cell HMRs. (B) This graph shows S-LDSC results for alanine transaminase. The data includes the annotations from (A) in addition to developmentally derived Liver HMRs.

**S1 Table.** Inter-HMR lengths by cell type. The table shows the distance between HMRs per cell type. Values represent the inter-HMR distances at different quantiles. The values have also been calculated at the .45 and .55 quantile values for more specificity around the median. Max values were limited to a maximum of 50kbp.

| Quantile | H1 ESC | Fetal Spinal | Fetal Heart | Adrenal | Liver | B cell |
| --- | --- | --- | --- | --- | --- | --- |
| 0.1 | 227 | 881 | 761 | 673 | 633 | 436.3 |
| 0.2 | 1283 | 1770 | 1626 | 1678 | 1508 | 1259 |
| 0.3 | 2953.8 | 2960 | 2787 | 2969 | 2716.8 | 2410 |
| 0.4 | 5626.6 | 4574 | 4329 | 4777 | 441.4 | 4030 |
| 0.45 | 7102.65 | 5484 | 5309 | 5990 | 5540 | 4998 |
| 0.5 | 8921 | 6555 | 6361 | 7311 | 6846 | 6142 |
| 0.55 | 11013.15 | 7805 | 7689 | 8799.3 | 8461 | 7474 |
| 0.6 | 13299.8 | 9237 | 9183 | 10524 | 10148.2 | 9112.8 |
| 0.7 | 18970 | 13267 | 13231 | 15015.6 | 14803 | 13573.1 |
| 0.8 | 26779 | 19213 | 19414 | 21739.2 | 21582 | 19841.2 |
| 0.9 | 37041.2 | 29618 | 29903 | 32016.2 | 31880.6 | 30592.7 |

**S2 Table.**
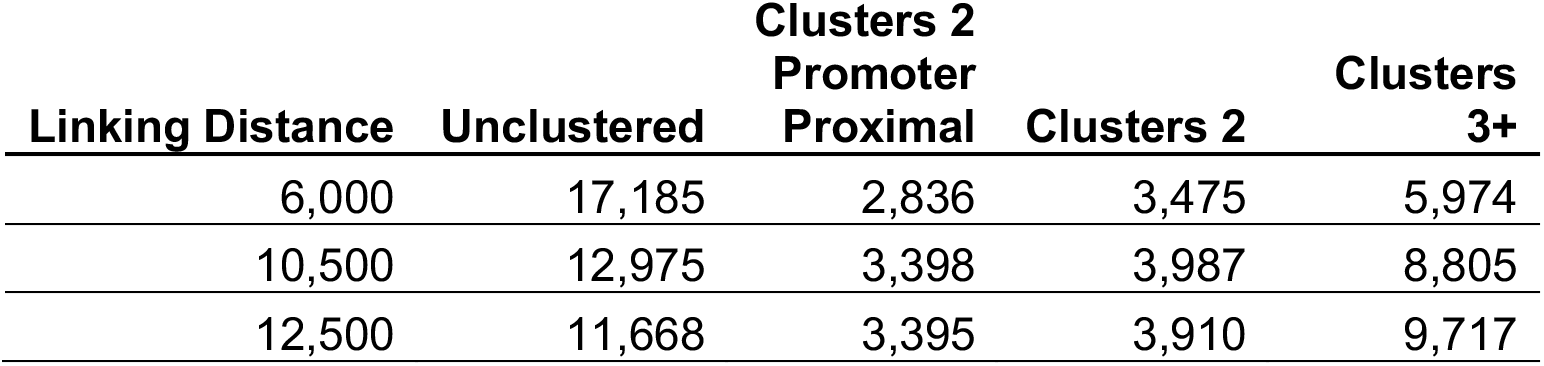
Clustering group counts by clustering distance. Table of HMR counts for clustering groups: unclustered; clusters of 2 HMRs where one is a RefSeq promoter; clusters of 2 non-coding HMRs; and clusters of 3+ HMRs. Linking distances include 6kb (used for our analyses); 10.5kb (approximate mean inter-HMR length); and 12.5kb (distance commonly used in super-enhancer definitions applied to histone modification ChIP-seq datasets).

**S3 Table.** List of 79 summary statistics files used for S-LDSC analyses. Table of trait names used in S-LDSC. Files were sourced from the Alkes Price lab website or Neale lab LDSC summary statistics browser.

**Traits:**
|  |  |
| --- | --- |
| Albumin | I9_IHD_wideDefinition |
| ALP | ICD10_I42_Cardiomyopathy |
| ALT | ICD10_I48_atrialFibrillationAndFlutter |
| Angina_byDoctor | IGF1 |
| Apolipoprotein_B | Liver_chirrosis |
| AST | lung_FEV1FVCzSMOKE |
| blood_EOSINOPHIL_COUNT | lung_FVCzSMOKE |
| blood_PLATELET_COUNT | Lymphocyte_count |
| blood_RBC_DISTRIB_WIDTH | LymphoidLeukemia |
| blood_RED_COUNT | mental_NEUROTICISM |
| blood_WHITE_COUNT | Myocardial_infarction |
| bmd_HEEL_TSCOREz | Neutrophil_count |
| body_BALDING1 | other_MORNINGPERSON |
| body_BMIz | PASS_AgeFirstBirth |
| body_HEIGHTz | PASS_Anorexia |
| body_WHRadjBMIz | PASS_Autism |
| bp_SYSTOLICadjMEDz | PASS_BMI1 |
| Cadiomyopathy_andOther | PASS_Coronary_Artery_Disease |
| CardiacArrythm | PASS_Crohns_Disease |
| Cholesterol | PASS_DS |
| Coffee_type | PASS_Ever_Smoked |
| Congen_Heart_andGreatArteries | PASS_HDL |
| cov_EDU_YEARS | PASS_Height1 |
| cov_SMOKING_STATUS | PASS_LDL |
| disease_AID_SURE | PASS_NumberChildrenEverBorn |
| disease_ALLERGY_ECZEMA_DIAGNOSED | PASS_Rheumatoid_Arthritis |
| disease_DERMATOLOGY | PASS_Schizophrenia |
| disease_HI_CHOL_SELF_REP | PASS_Type_2_Diabetes |
| disease_HYPOTHYROIDISM_SELF_REP | PASS_Ulcerative_Colitis |
| disease_RESPIRATORY_ENT | PASS_Years_of_Education2 |
| Diseases_of_liver | pigment_HAIR |
| disease_T2D | pigment_SKIN |
| ECG_load | pigment_SUNBURN |
| ECG_phaseTime | pigment_TANNING |
| ECG | Primary_lymphoid_neoplasms |
| Haematocrit_percentage | repro_MENARCHE_AGE |
| HeartAttack_byDoctor | repro_MENOPAUSE_AGE |
| HighBloodPressure_byDoctor | RheumatoidFactor |
| I25_chronicIHD | Triglycerides |
| I9_Cardiomyopathy |  |

## REFERENCES

1. Van der Ploeg L, Flavell R. DNA methylation in the human γδβ-globin locus in erythroid and nonerythroid tissues. Cell. 1980;19(4):947–58.

2. Gruenbaum Y, Stein R, Cedar H, Razin A. Methylation of CpG sequences in eukaryotic DNA. FEBS letters. 1981;124(1):67–71.

3. Kunnath L, Locker J. Characterization of DNA methylation in the rat. Biochimica et Biophysica Acta (BBA)-Gene Structure and Expression. 1982;699(3):264–71.

4. Lister R, Pelizzola M, Dowen RH, Hawkins RD, Hon G, Tonti-Filippini J, et al. Human DNA methylomes at base resolution show widespread epigenomic differences. Nature. 2009;462(7271):315–22.

5. Gu J, Stevens M, Xing X, Li D, Zhang B, Payton JE, et al. Mapping of variable DNA methylation across multiple cell types defines a dynamic regulatory landscape of the human genome. G3: Genes, Genomes, Genetics. 2016;6(4):973–86.

6. Gama-Sosa MA, Midgett RM, Slagel VA, Githens S, Kuo KC, Gehrke CW, et al. Tissue-specific differences in DNA methylation in various mammals. Biochimica et Biophysica Acta (BBA)-Gene Structure and Expression. 1983;740(2):212–9.

7. Meissner A, Mikkelsen TS, Gu H, Wernig M, Hanna J, Sivachenko A, et al. Genome-scale DNA methylation maps of pluripotent and differentiated cells. Nature. 2008;454(7205):766–70.

8. Portela A, Esteller M. Epigenetic modifications and human disease. Nature biotechnology. 2010;28(10):1057–68.

9. Hodges E, Molaro A, Dos Santos CO, Thekkat P, Song Q, Uren PJ, et al. Directional DNA methylation changes and complex intermediate states accompany lineage specificity in the adult hematopoietic compartment. Molecular cell. 2011;44(1):17–28.

10. Molaro A, Hodges E, Fang F, Song Q, McCombie WR, Hannon GJ, et al. Sperm methylation profiles reveal features of epigenetic inheritance and evolution in primates. Cell. 2011;146(6):1029–41.

11. Bock C, Beerman I, Lien W-H, Smith ZD, Gu H, Boyle P, et al. DNA methylation dynamics during in vivo differentiation of blood and skin stem cells. Molecular cell. 2012;47(4):633–47.

12. Barnett KR, Decato BE, Scott TJ, Hansen TJ, Chen B, Attalla J, et al. ATAC-Me captures prolonged DNA methylation of dynamic chromatin accessibility loci during cell fate transitions. Molecular cell. 2020;77(6):1350–64. e6.

13. He Y, Hariharan M, Gorkin DU, Dickel DE, Luo C, Castanon RG, et al. Spatiotemporal DNA methylome dynamics of the developing mouse fetus. Nature. 2020;583(7818):752–9.

14. Razin A, Szyf M. DNA methylation patterns formation and function. Biochimica et Biophysica Acta (BBA)-Gene Structure and Expression. 1984;782(4):331–42.

15. Weber M, Hellmann I, Stadler MB, Ramos L, Pääbo S, Rebhan M, et al. Distribution, silencing potential and evolutionary impact of promoter DNA methylation in the human genome. Nature genetics. 2007;39(4):457–66.

16. Eckhardt F, Lewin J, Cortese R, Rakyan VK, Attwood J, Burger M, et al. DNA methylation profiling of human chromosomes 6, 20 and 22. Nature genetics. 2006;38(12):1378–85.

17. Neri F, Rapelli S, Krepelova A, Incarnato D, Parlato C, Basile G, et al. Intragenic DNA methylation prevents spurious transcription initiation. Nature. 2017;543(7643):72–7.

18. Kundaje A, Meuleman W, Ernst J, Bilenky M, Yen A, Heravi-Moussavi A, et al. Integrative analysis of 111 reference human epigenomes. Nature. 2015;518(7539):317–30.

19. Schlesinger F, Smith AD, Gingeras TR, Hannon GJ, Hodges E. De novo DNA demethylation and noncoding transcription define active intergenic regulatory elements. Genome research. 2013;23(10):1601–14.

20. Schultz MD, He Y, Whitaker JW, Hariharan M, Mukamel EA, Leung D, et al. Human body epigenome maps reveal noncanonical DNA methylation variation. Nature. 2015;523(7559):212–6.

21. Stadler MB, Murr R, Burger L, Ivanek R, Lienert F, Scholer A, et al. DNA-binding factors shape the mouse methylome at distal regulatory regions. Nature. 2011;480(7378):490–5.

22. Ziller MJ, Gu H, Müller F, Donaghey J, Tsai LT-Y, Kohlbacher O, et al. Charting a dynamic DNA methylation landscape of the human genome. Nature. 2013;500(7463):477–81.

23. Mendizabal I, Shi L, Keller TE, Konopka G, Preuss TM, Hsieh T-F, et al. Comparative methylome analyses identify epigenetic regulatory loci of human brain evolution. Molecular biology and evolution. 2016;33(11):2947–59.

24. Yuan W, Xia Y, Bell CG, Yet I, Ferreira T, Ward KJ, et al. An integrated epigenomic analysis for type 2 diabetes susceptibility loci in monozygotic twins. Nature communications. 2014;5(1):1–7.

25. Hindorff LA, Sethupathy P, Junkins HA, Ramos EM, Mehta JP, Collins FS, et al. Potential etiologic and functional implications of genome-wide association loci for human diseases and traits. Proceedings of the National Academy of Sciences. 2009;106(23):9362–7.

26. Li Y, Rivera CM, Ishii H, Jin F, Selvaraj S, Lee AY, et al. CRISPR reveals a distal super-enhancer required for Sox2 expression in mouse embryonic stem cells. PloS one. 2014;9(12):e114485.

27. Hon GC, Rajagopal N, Shen Y, McCleary DF, Yue F, Dang MD, et al. Epigenetic memory at embryonic enhancers identified in DNA methylation maps from adult mouse tissues. Nature genetics. 2013;45(10):1198–206.

28. Bell E, Curry EW, Megchelenbrink W, Jouneau L, Brochard V, Tomaz RA, et al. Dynamic CpG methylation delineates subregions within super-enhancers selectively decommissioned at the exit from naive pluripotency. Nature communications. 2020;11(1):1–16.

29. Molaro A, Falciatori I, Hodges E, Aravin AA, Marran K, Rafii S, et al. Two waves of de novo methylation during mouse germ cell development. Genes & development. 2014;28(14):1544–9.

30. Song Q, Decato B, Hong EE, Zhou M, Fang F, Qu J, et al. A reference methylome database and analysis pipeline to facilitate integrative and comparative epigenomics. PloS one. 2013;8(12):e81148.

31. Phillips JE, Corces VG. CTCF: master weaver of the genome. Cell. 2009;137(7):1194–211.

32. Rodda DJ, Chew J-L, Lim L-H, Loh Y-H, Wang B, Ng H-H, et al. Transcriptional regulation of nanog by OCT4 and SOX2. Journal of Biological Chemistry. 2005;280(26):24731–7.

33. Kistler B, Pfisterer P, Wirth T. Lymphoid-and myeloid-specific activity of the PU. 1 promoter is determined by the combinatorial action of octamer and ets transcription factors. Oncogene. 1995;11(6):1095–106.

34. Chen J. The proto-oncogene c-ets is preferentially expressed in lymphoid cells. Molecular and cellular biology. 1985;5(11):2993–3000.

35. Åkerblad P, Sigvardsson M. Early B cell factor is an activator of the B lymphoid kinase promoter in early B cell development. The Journal of Immunology. 1999;163(10):5453–61.

36. McLean CY, Bristor D, Hiller M, Clarke SL, Schaar BT, Lowe CB, et al. GREAT improves functional interpretation of cis-regulatory regions. Nature biotechnology. 2010;28(5):495–501.

37. Ji H, Ehrlich LI, Seita J, Murakami P, Doi A, Lindau P, et al. Comprehensive methylome map of lineage commitment from haematopoietic progenitors. Nature. 2010;467(7313):338–42.

38. Bröske A-M, Vockentanz L, Kharazi S, Huska MR, Mancini E, Scheller M, et al. DNA methylation protects hematopoietic stem cell multipotency from myeloerythroid restriction. Nature genetics. 2009;41(11):1207–15.

39. Hendriks J, Gravestein LA, Tesselaar K, van Lier RA, Schumacher TN, Borst J. CD27 is required for generation and long-term maintenance of T cell immunity. Nature immunology. 2000;1(5):433–40.

40. Agematsu K, Hokibara S, Nagumo H, Komiyama A. CD27: a memory B-cell marker. Immunology today. 2000;21(5):204–6.

41. Lens SM, Tesselaar K, van Oers MH, van Lier RA, editors. Control of lymphocyte function through CD27–CD70 interactions. Seminars in immunology; 1998: Elsevier.

42. Whyte WA, Orlando DA, Hnisz D, Abraham BJ, Lin CY, Kagey MH, et al. Master transcription factors and mediator establish super-enhancers at key cell identity genes. Cell. 2013;153(2):307–19.

43. Hnisz D, Abraham BJ, Lee TI, Lau A, Saint-André V, Sigova AA, et al. Super-enhancers in the control of cell identity and disease. Cell. 2013;155(4):934–47.

44. Lovén J, Hoke HA, Lin CY, Lau A, Orlando DA, Vakoc CR, et al. Selective inhibition of tumor oncogenes by disruption of super-enhancers. Cell. 2013;153(2):320–34.

45. Sabari BR, Dall’Agnese A, Boija A, Klein IA, Coffey EL, Shrinivas K, et al. Coactivator condensation at super-enhancers links phase separation and gene control. Science. 2018;361(6400):eaar3958.

46. Vahedi G, Kanno Y, Furumoto Y, Jiang K, Parker SC, Erdos MR, et al. Super-enhancers delineate disease-associated regulatory nodes in T cells. Nature. 2015;520(7548):558–62.

47. Parker SC, Stitzel ML, Taylor DL, Orozco JM, Erdos MR, Akiyama JA, et al. Chromatin stretch enhancer states drive cell-specific gene regulation and harbor human disease risk variants. Proceedings of the National Academy of Sciences. 2013;110(44):17921–6.

48. Ernst J, Kellis M. ChromHMM: automating chromatin-state discovery and characterization. Nature methods. 2012;9(3):215–6.

49. Ernst J, Kellis M. Chromatin-state discovery and genome annotation with ChromHMM. Nature protocols. 2017;12(12):2478–92.

50. Heintzman ND, Stuart RK, Hon G, Fu Y, Ching CW, Hawkins RD, et al. Distinct and predictive chromatin signatures of transcriptional promoters and enhancers in the human genome. Nature genetics. 2007;39(3):311–8.

51. Creyghton MP, Cheng AW, Welstead GG, Kooistra T, Carey BW, Steine EJ, et al. Histone H3K27ac separates active from poised enhancers and predicts developmental state. Proceedings of the National Academy of Sciences. 2010;107(50):21931–6.

52. Visel A, Blow MJ, Li Z, Zhang T, Akiyama JA, Holt A, et al. ChIP-seq accurately predicts tissue-specific activity of enhancers. Nature. 2009;457(7231):854–8.

53. Capra JA. Extrapolating histone marks across developmental stages, tissues, and species: an enhancer prediction case study. BMC genomics. 2015;16(1):1–9.

54. Huang J, Li K, Cai W, Liu X, Zhang Y, Orkin SH, et al. Dissecting super-enhancer hierarchy based on chromatin interactions. Nature communications. 2018;9(1):1–12.

55. Fraser P, Pruzina S, Antoniou M, Grosveld F. Each hypersensitive site of the human beta-globin locus control region confers a different developmental pattern of expression on the globin genes. Genes & development. 1993;7(1):106–13.

56. Hansen TJ, Hodges E. ATAC-STARR-seq reveals transcription factor–bound activators and silencers within chromatin-accessible regions of the human genome. Genome Research. 2022;32(8):1529–41.

57. Wang X, He L, Goggin SM, Saadat A, Wang L, Sinnott-Armstrong N, et al. High-resolution genome-wide functional dissection of transcriptional regulatory regions and nucleotides in human. Nature communications. 2018;9(1):1–15.

58. Fulco CP, Nasser J, Jones TR, Munson G, Bergman DT, Subramanian V, et al. Activity-by-contact model of enhancer–promoter regulation from thousands of CRISPR perturbations. Nature genetics. 2019;51(12):1664–9.

59. Consortium EP. A user’s guide to the encyclopedia of DNA elements (ENCODE). PLoS biology. 2011;9(4):e1001046.

60. Genomes Project C, Abecasis GR, Auton A, Brooks LD, DePristo MA, Durbin RM, et al. An integrated map of genetic variation from 1,092 human genomes. Nature. 2012;491(7422):56–65.

61. Roederer M, Quaye L, Mangino M, Beddall MH, Mahnke Y, Chattopadhyay P, et al. The genetic architecture of the human immune system: a bioresource for autoimmunity and disease pathogenesis. Cell. 2015;161(2):387–403.

62. Chun S, Casparino A, Patsopoulos NA, Croteau-Chonka DC, Raby BA, De Jager PL, et al. Limited statistical evidence for shared genetic effects of eQTLs and autoimmune-disease-associated loci in three major immune-cell types. Nat Genet. 2017.

63. Guo MH, Nandakumar SK, Ulirsch JC, Zekavat SM, Buenrostro JD, Natarajan P, et al. Comprehensive population-based genome sequencing provides insight into hematopoietic regulatory mechanisms. Proc Natl Acad Sci U S A. 2017;114(3):E327–E36.

64. Vockley CM, Barrera A, Reddy TE. Decoding the role of regulatory element polymorphisms in complex disease. Curr Opin Genet Dev. 2017;43:38–45.

65. Maurano MT, Humbert R, Rynes E, Thurman RE, Haugen E, Wang H, et al. Systematic localization of common disease-associated variation in regulatory DNA. Science. 2012;337(6099):1190–5.

66. Koues OI, Kowalewski RA, Chang LW, Pyfrom SC, Schmidt JA, Luo H, et al. Enhancer sequence variants and transcription-factor deregulation synergize to construct pathogenic regulatory circuits in B-cell lymphoma. Immunity. 2015;42(1):186–98.

67. Chen L, Ge B, Casale FP, Vasquez L, Kwan T, Garrido-Martin D, et al. Genetic Drivers of Epigenetic and Transcriptional Variation in Human Immune Cells. Cell. 2016;167(5):1398–414.e24.

68. Farh KK, Marson A, Zhu J, Kleinewietfeld M, Housley WJ, Beik S, et al. Genetic and epigenetic fine mapping of causal autoimmune disease variants. Nature. 2015;518(7539):337–43.

69. Finucane HK, Bulik-Sullivan B, Gusev A, Trynka G, Reshef Y, Loh PR, et al. Partitioning heritability by functional annotation using genome-wide association summary statistics. Nat Genet. 2015;47(11):1228–35.

70. Zhang B, Kirov S, Snoddy J. WebGestalt: an integrated system for exploring gene sets in various biological contexts. Nucleic acids research. 2005;33(suppl_2):W741–W8.

71. Liao Y, Wang J, Jaehnig EJ, Shi Z, Zhang B. WebGestalt 2019: gene set analysis toolkit with revamped UIs and APIs. Nucleic acids research. 2019;47(W1):W199–W205.

72. Kuroda A, Rauch TA, Todorov I, Ku HT, Al-Abdullah IH, Kandeel F, et al. Insulin gene expression is regulated by DNA methylation. PloS one. 2009;4(9):e6953.

73. Schmidl C, Klug M, Boeld TJ, Andreesen R, Hoffmann P, Edinger M, et al. Lineage-specific DNA methylation in T cells correlates with histone methylation and enhancer activity. Genome research. 2009;19(7):1165–74.

74. Bloushtain-Qimron N, Yao J, Snyder EL, Shipitsin M, Campbell LL, Mani SA, et al. Cell type-specific DNA methylation patterns in the human breast. Proceedings of the National Academy of Sciences. 2008;105(37):14076–81.

75. Deaton AM, Webb S, Kerr AR, Illingworth RS, Guy J, Andrews R, et al. Cell type–specific DNA methylation at intragenic CpG islands in the immune system. Genome research. 2011;21(7):1074–86.

76. Park I-H, Wen B, Murakami P, Aryee MJ, Irizarry R, Herb B, et al. Differential methylation of tissue-and cancer-specific CpG island shores distinguishes human induced pluripotent stem cells, embryonic stem cells and fibroblasts. Nature genetics. 2009;41(12): 1350–3.

77. Frigola J, Song J, Stirzaker C, Hinshelwood RA, Peinado MA, Clark SJ. Epigenetic remodeling in colorectal cancer results in coordinate gene suppression across an entire chromosome band. Nature genetics. 2006;38(5):540–9.

78. Irizarry RA, Ladd-Acosta C, Wen B, Wu Z, Montano C, Onyango P, et al. The human colon cancer methylome shows similar hypo-and hypermethylation at conserved tissue-specific CpG island shores. Nature genetics. 2009;41(2):178–86.

79. Bell JT, Tsai P-C, Yang T-P, Pidsley R, Nisbet J, Glass D, et al. Epigenome-wide scans identify differentially methylated regions for age and age-related phenotypes in a healthy ageing population. PLoS genetics. 2012;8(4):e1002629.

80. Hotta K, Kitamoto A, Kitamoto T, Ogawa Y, Honda Y, Kessoku T, et al. Identification of differentially methylated region (DMR) networks associated with progression of nonalcoholic fatty liver disease. Scientific reports. 2018;8(1):1–11.

81. Schulz M, Teissandier A, de la Mata E, Armand M, Iranzo J, El Marjou F, et al. DNA methylation restricts coordinated germline and neural fates in embryonic stem cell differentiation. bioRxiv. 2022:2022.10.22.513040.

82. Guerin LN, Barnett KR, Hodges E. Dual detection of chromatin accessibility and DNA methylation using ATAC-Me. Nature Protocols. 2021;16(12):5377–97.

83. Karolchik D, Baertsch R, Diekhans M, Furey TS, Hinrichs A, Lu Y, et al. The UCSC genome browser database. Nucleic acids research. 2003;31(1):51–4.

84. Quinlan AR, Hall IM. BEDTools: a flexible suite of utilities for comparing genomic features. Bioinformatics. 2010;26(6):841–2.

85. Kolde R. Pheatmap: pretty heatmaps. R package version. 2012;1(2):726.

86. Heinz S, Benner C, Spann N, Bertolino E, Lin YC, Laslo P, et al. Simple combinations of lineage-determining transcription factors prime cis-regulatory elements required for macrophage and B cell identities. Molecular cell. 2010;38(4):576–89.

87. Wickham H. Package ‘ggplot2’: elegant graphics for data analysis. Springer-Verlag New York doi. 2016;10:978–0.

88. Jourquin J, Duncan D, Shi Z, Zhang B. GLAD4U: deriving and prioritizing gene lists from PubMed literature. BMC genomics. 2012;13(8):1–12.

89. Phanstiel DH, Boyle AP, Araya CL, Snyder MP. Sushi. R: flexible, quantitative and integrative genomic visualizations for publication-quality multi-panel figures. Bioinformatics. 2014;30(19):2808–10.

